# Evolutionary constraints limit additional mammalian adaptation of bovine-derived H5N1 influenza viruses in ferrets

**DOI:** 10.64898/2026.07.20.739655

**Authors:** Heather M. Machkovech, Wanting Wei, Chunyang Gu, Lizheng Guan, Asim Biswas, Tadashi Maemura, Tong Wang, Lavanya Babujee, Peter J. Halfmann, Amie J. Eisfeld, Gabriele Neumann, Yoshihiro Kawaoka, Katia Koelle, Thomas C. Friedrich

## Abstract

The outbreak of clade 2.3.4.4b H5N1 viruses among U.S. dairy cattle has raised concerns that sustained circulation among agricultural mammals could facilitate viral adaptation toward efficient human transmission. However, the evolutionary dynamics governing such adaptation remain poorly understood. Here we investigated the evolution of two bovine-derived H5N1 B3.13 genotype viruses during infection and airborne transmission in ferrets, building on prior characterization of their robust replication and inefficient airborne transmission. Within hosts, viral genetic diversity was limited and viruses were subject to genetic drift and weak purifying selection. Transmission, when it occurred, was characterized by stringent bottlenecks that sharply reduced viral genetic diversity. We found no evidence of mammalian adaptation during infection or transmission. Together, these findings indicate that bovine-derived H5N1 viruses face evolutionary constraints during acute mammalian infection and transmission, limiting movement toward enhanced airborne spread. These constraints may help explain why efficient mammalian replication does not necessarily coincide with efficient transmission. Continued circulation of HA clade 2.3.4.4b viruses nevertheless creates repeated opportunities for rare but consequential evolutionary events, underscoring the importance of sustained surveillance and risk mitigation.

## Introduction

In early 2024, clade 2.3.4.4b H5N1 was first detected in dairy cattle in Texas, and since then more than 1,000 infected herds have been reported across multiple U.S. states ^1^.

The vast majority of dairy herd infections have involved viruses of the B3.13 genotype, which emerged following an initial wild-bird-to-cattle spillover event, but H5N1 viruses of genotype D1.1 have also been detected in Arizona, Nevada, and Wisconsin dairy cattle, representing three additional independent introductions from wild birds ^2^. Infection in dairy cattle is characterized primarily by mastitis, resulting in exceptionally high infectious titers in raw milk ^3,4^. During April 1, 2024–June 30, 2025, 70 human H5N1 infections were reported in the United States, the majority associated with occupational exposure to infected cattle or poultry, including one fatal case ^5^. Clinical disease has generally been mild, dominated by conjunctivitis with limited respiratory involvement ^6,7^ and no sustained human-to-human transmission has been observed. Nevertheless, the scale and persistence of this outbreak raise concern that prolonged circulation in mammals could facilitate viral adaptation with implications for zoonotic and pandemic risk.

Studies using ferret models, a well-established system for assessing influenza virus transmissibility and pathogenesis, have demonstrated that cattle-outbreak-associated clade 2.3.4.4b H5N1 viruses can replicate efficiently in the ferret respiratory tract and, in some settings, produce infectious virus in exposed animals or in air samples 8–10.

For the two viruses analyzed here, A/dairy cattle/New Mexico/A240920343-93/2024 (NM93-H5N1, derived from milk from an infected cow) replicated efficiently in inoculated ferrets but produced no detectable infectious virus in exposed recipients, although one recipient seroconverted ^10^. A/Texas/37/2024 (huTX37-H5N1, derived from a clinical isolate from a patient with conjunctivitis) likewise replicated efficiently in inoculated ferrets, but produced detectable infection in only a subset of exposed recipients ^9^. These findings indicate that robust replication in a mammalian host does not necessarily predict efficient productive airborne transmission.

Hemagglutinin (HA) receptor binding specificity is crucial in determining the range of host species in which influenza A viruses can efficiently transmit ^11^. The upper respiratory tract of mammals, including ferrets and humans, is dominated by α2,6-linked sialic acid receptors, while avian influenza viruses typically preferentially bind α2,3-linked sialic acids ^12^. Shifts toward α2,6 receptor usage appear to be required for efficient airborne transmission in mammals ^11^ and, in some genetic backgrounds, including bovine-derived clade 2.3.4.4b H5 hemagglutinins, can be conferred by single amino acid substitutions in HA ^13^. Prior work from our group and others has shown that avian-like influenza viruses, including recombinant 1918-like H1N1 and H5N1 viruses, can undergo rapid adaptive evolution in HA during replication and transmission in ferrets ^14,15^. Previous studies have shown that NM93-H5N1 and huTX37-H5N1 primarily bind α2,3-linked sialic acids, although some evidence suggests limited binding to α2,6-linked sialylglycopolymers ^9,10,16^. Because huTX37-H5N1 produced detectable infection in a subset of airborne-exposed recipient ferrets, these viruses provide an opportunity to ask whether the combination of diversifying selection within hosts and selective bottlenecks during mammalian transmission could drive the emergence of enhanced mammalian adaptation in HA.

Despite growing phenotypic data on replication, pathogenicity, and transmissibility, the evolutionary processes governing adaptation of bovine-derived H5N1 viruses during mammalian infection and transmission remain poorly understood. Here, we address these questions by analyzing within-host and between-host evolutionary dynamics of NM93-H5N1 and huTX37-H5N1 using deep sequencing data generated from previously described ferret infection and airborne transmission experiments ^9,10^. By quantifying intrahost diversity, selection pressures, and transmission bottleneck sizes, we aim to define the constraints and contingencies that govern mammalian adaptation of bovine-derived H5N1 viruses and to clarify how replication fitness, transmission efficiency, and evolutionary potential are related in this emerging zoonotic system.

## Results

### Viral stocks differ at known mammalian-adaptive sites

To contextualize subsequent evolutionary analyses, we first compared the consensus sequences of the NM93-H5N1 and huTX37-H5N1 viral stocks used in these experiments (Fig. 1). The two viruses are closely related, differing at only 12 amino acid positions across the genome. Notably, both virus isolates harbor distinct polymerase mutations previously associated with mammalian adaptation. NM93-H5N1 contains PB2 M631L, a mutation that is fixed among circulating bovine H5N1 viruses and has been shown to enhance interactions between the viral polymerase complex and mammalian ANP32 proteins ^9,17–20^. In contrast, huTX37-H5N1 harbors PB2 E627K, a well-established mammalian-adaptive marker that confers a greater increase in polymerase activity than PB2 M631L^21,22^.

**Figure 1.**
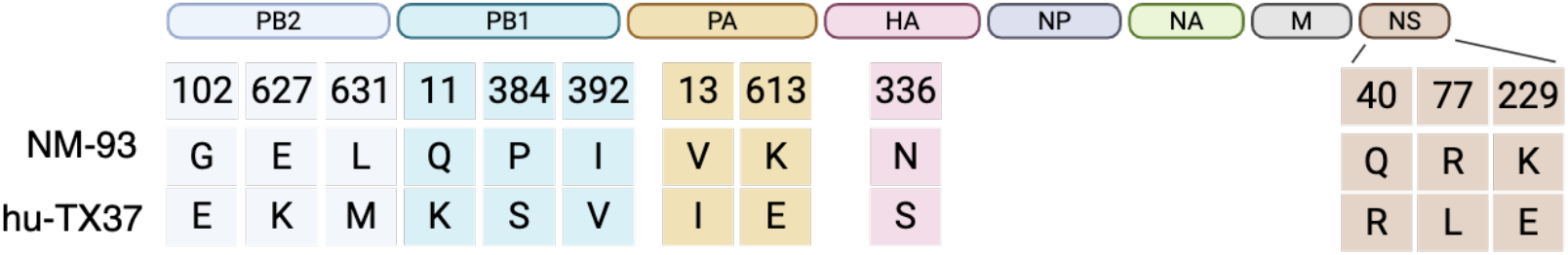
Amino acid differences between NM93-H5N1 and huTX37-H5N1, the two virus isolates used in the study.

### Summary of infection/transmission experiments

Here we examined nasal wash specimens from previously described ferret infection and airborne transmission experiments involving NM93-H5N1 and huTX37-H5N1 (Fig. 2). In the NM93-H5N1 study, donor ferrets were inoculated with 10⁶ PFU and each was paired with a naïve recipient (n = 4 pairs). While NM93-H5N1 replicated efficiently in donor animals, no infectious virus was detected in recipient ferrets although one recipient seroconverted, indicating exposure in the absence of detectable productive infection. In comparison, in the same experimental setup, the seasonal human influenza virus A/Isumi/UT-KK001-01/2018 (H1N1) transmitted efficiently to all exposed recipients. In a separate study, huTX37-H5N1 was evaluated across a range of inoculation doses (10⁶, 10³, 10², or 10¹ PFU; n = 6 ferrets per dose) and transmitted inefficiently, with infectious virus detected in 5 of 24 exposed recipients across dose groups; an additional exposed recipient seroconverted without detectable infectious virus. All ferrets infected with huTX37-H5N1, including the 5 exposed recipients who became infected, succumbed to infection within 7 days.

**Figure 2.**
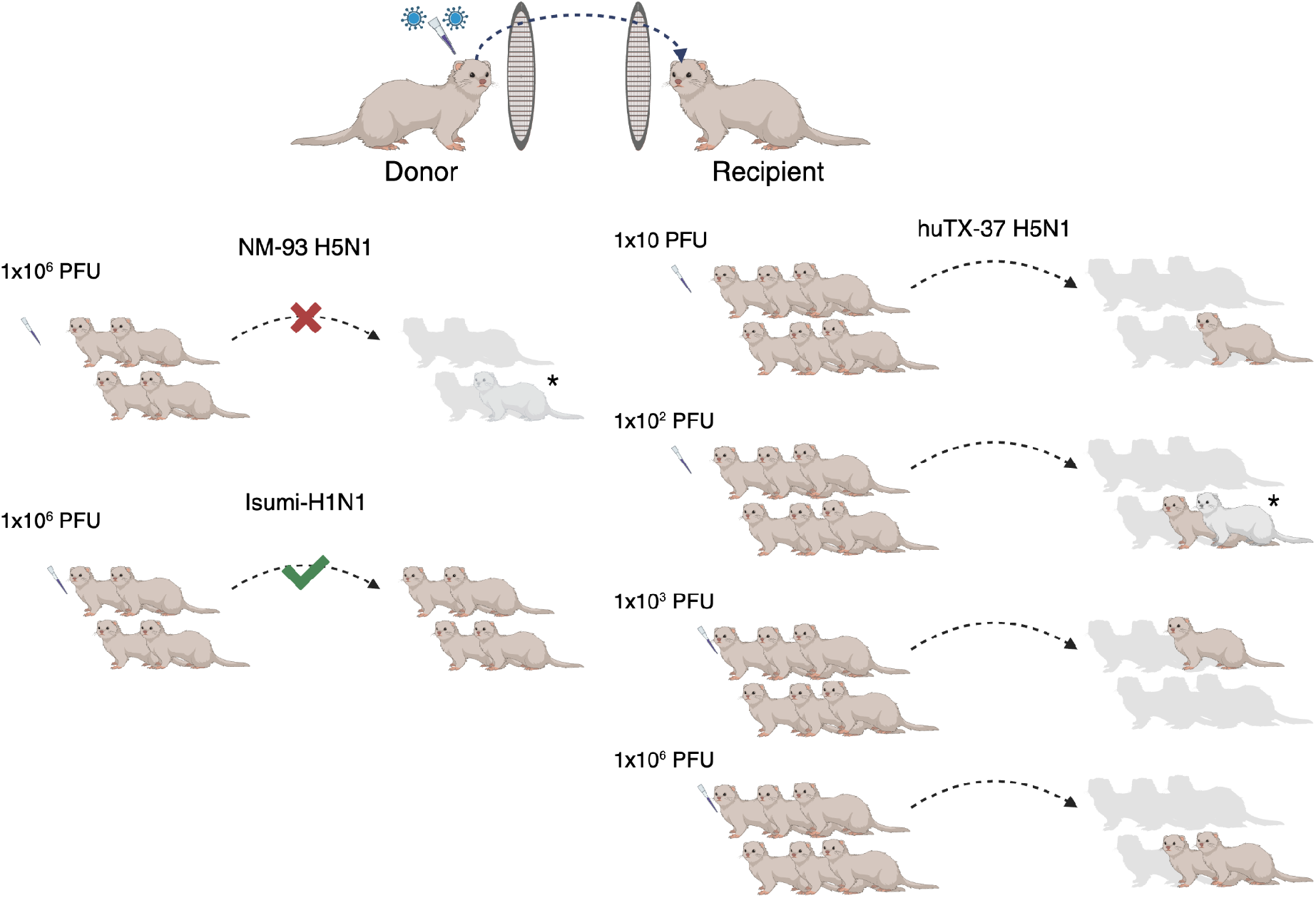
Overview of ferret transmission experiments. In the first study, donor ferrets were inoculated with the bovine isolate NM93-H5N1 or the human seasonal virus A/Isumi/UT-KK001-01/2018 (Isumi-H1N1) and paired with naive recipients (n=4 pairs per virus; left). In the second experiment, donor ferrets were inoculated with the human H5N1 isolate huTX37-H5N1 at 4 different doses and then paired with naive recipients (n=6 pairs per dose; right). Animals with detectable virus are colored in tan. Recipient animals that seroconverted but did not have detectable virus are outlined in gray and denoted with an asterisk.

Together, these experiments generated a defined set of donor samples from both viruses and recipient samples from huTX37-H5N1 productive airborne transmission events. Nasal wash samples with detectable infectious virus were subjected to deep sequencing; sequence data for huTX37-H5N1 were obtained from previously published data ^9^, whereas NM93-H5N1 samples were newly sequenced for this study.

### Limited within-host viral genetic diversity during ferret infection

We first characterized intra-sample single-nucleotide variants (iSNVs) present in the virus stocks used to inoculate ferrets. iSNVs were called using a previously described workflow, with a minimum frequency threshold of 3% and a minimum read depth of 200× ^23,24^. In the NM93- H5N1 inoculum, four iSNVs were detected, all below 10% frequency. Only one of these variants was nonsynonymous (PB1 P68T, 7.9%). Similarly, the huTX37-H5N1 inoculum contained five iSNVs, all below 20%, including three nonsynonymous mutations (PA L655F, NA G454D, and NS1 P164S) and two synonymous mutations (NA S405S and PB2 S688S). Thus, each viral inoculum contained only low levels of genetic diversity prior to infection.

We next assessed how viral populations diversified during infection by quantifying the number of iSNVs detected in each ferret over time. In NM93-H5N1–infected animals, iSNV counts increased modestly over the course of infection in two ferrets, while remaining relatively stable in the others (Fig. 3a). Notably, the highest iSNV counts in NM93-H5N1–infected animals occurred at day 7 post-infection, corresponding to the latest time point sampled, when viral loads had already declined to very low levels. iSNV counts remained largely unchanged across sampled time points in most huTX37-H5N1–infected ferrets (Fig. 3b). This pattern likely in part reflects the shorter duration of infection in huTX37-H5N1–infected donors: most succumbed by approximately day 5 post-infection, which limited later sampling and did not allow for quantification of within-host diversity beyond early infection.

**Figure 3.**
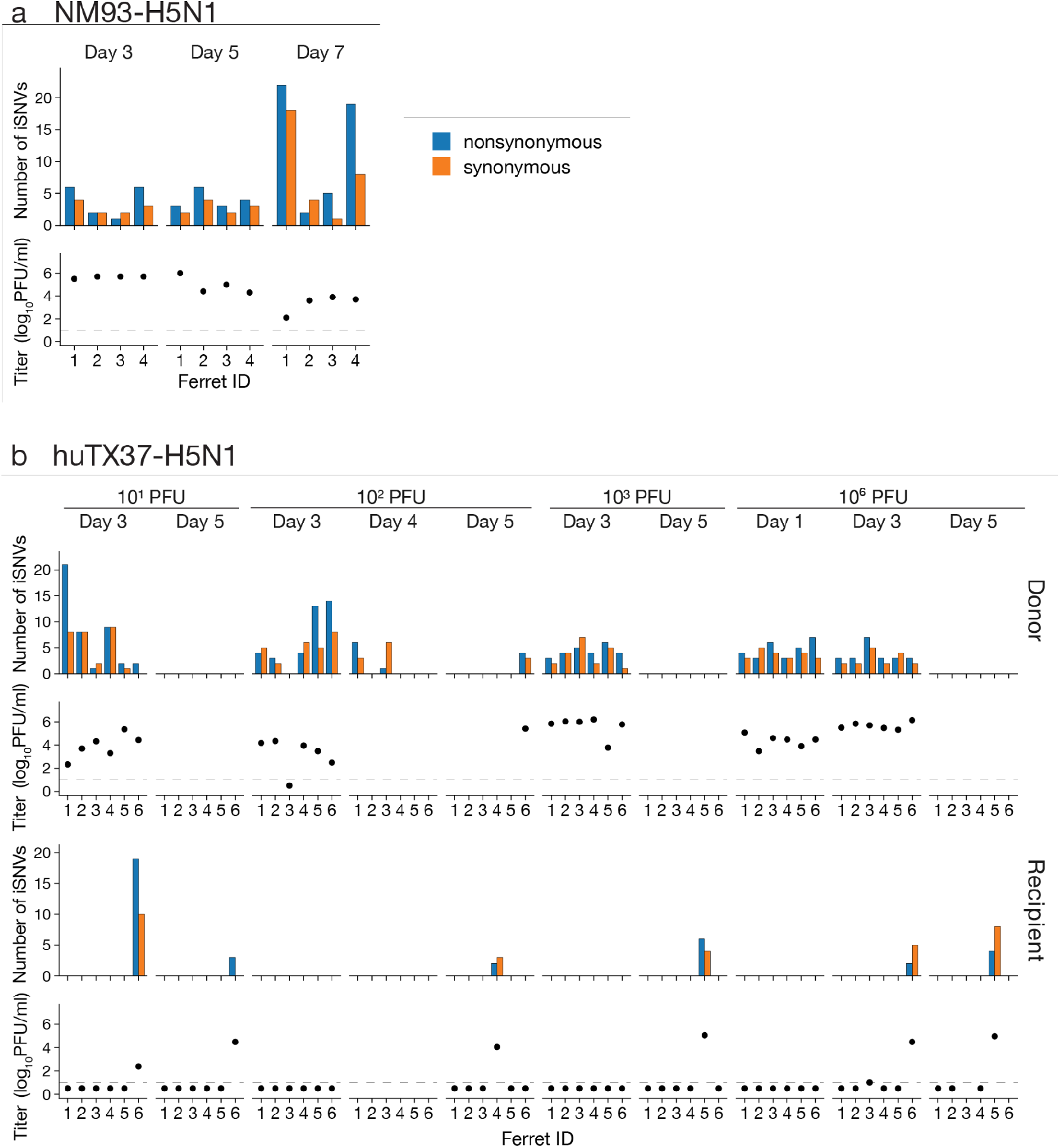
Number of synonymous and nonsynonymous iSNVs detected in each sample. (a) Number of iSNVs in the NM93-H5N1 study. (b) Number of iSNVs in the huTX37-H5N1 study, grouped by inoculation dose. Samples without bars do not have sequencing data. Viral titers are plotted below each sample. Dotted line denotes limit of detection.

Given the overall low level of within-host diversity, we next asked whether the detected variants included recurrent or high-frequency mutations that might indicate repeatable within-host adaptation. The genome positions of all detected iSNVs are shown in Supplemental Fig. 1. For visualization, only iSNVs detected in more than one sample and/or present at >20% frequency in at least one sample are displayed in Fig. 4. This analysis provides a summary of the major iSNVs observed during infection, while longitudinal allele-frequency dynamics are examined in the following section.

**Figure 4.**
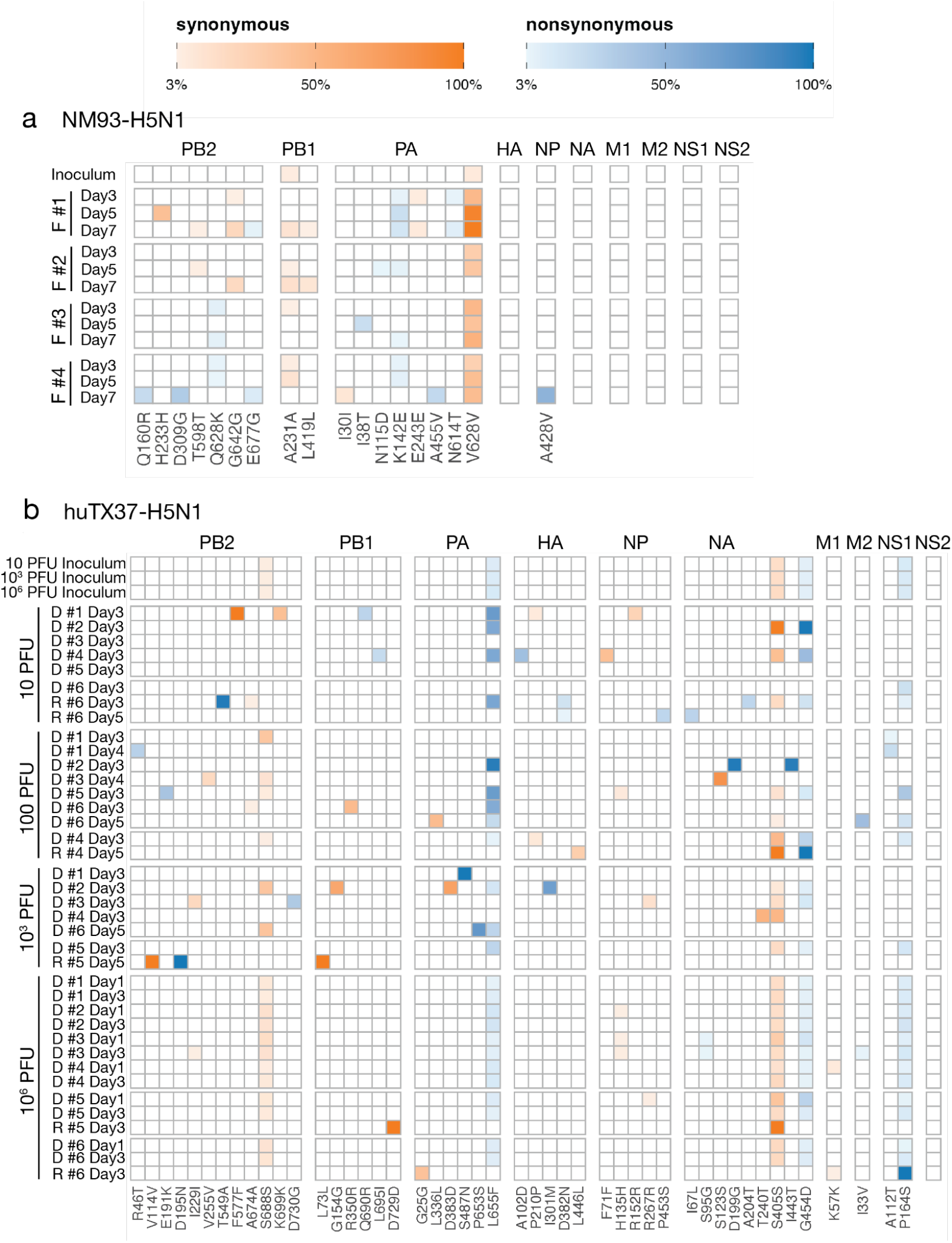
Heatmap of iSNV frequencies relative to the inoculum isolate across the influenza virus genome in infected ferrets. (a) Ferrets infected with NM93-H5N1. (b) Ferrets infected with huTX37-H5N1. Only iSNVs that occur in more than one sample or are present at >20% frequency in at least one sample are displayed. All iSNVs are displayed in Supplementary Fig.1.

Across the four NM93-H5N1–infected ferrets, only 17 mutations met this threshold. Of these 17, 15 occurred in the polymerase gene segments and 13 never exceeded a frequency of 30% (Fig. 4a). In addition, only two mutations reached majority frequency: NP A428V and PA V628V. PA K142E was detected in all four NM93-H5N1–infected ferrets at low frequency (<20%) across multiple time points. This variant was also detected in the inoculum below our stringent 3% iSNV calling threshold, suggesting that it was likely present as low-frequency standing variation in the inoculum. PA K142E has previously been associated with increased virulence of H5N1 viruses in mammalian models when combined with PB2 E627K; however, PB2 E627K was not detected in any NM93-H5N1 viruses in this study ^25^. PA K142E has also been detected sporadically among B3.13 cattle-lineage viruses. Another mutation of interest, PA I38T, was detected in a single NM93-H5N1–infected ferret at 5 days post-infection at approximately 20% frequency but was absent at earlier and later time points. Substitutions at PA position 38 are known to confer reduced susceptibility to baloxavir ^26^ and have been reported sporadically in avian influenza viruses, including B3.13 cattle-lineage viruses.

A similar overall pattern was observed in huTX37-H5N1–infected ferrets, although fewer longitudinal time points were available due to the high lethality of this virus. Across all donor and recipient huTX37-H5N1 samples, 230 unique iSNVs were detected, most of which remained at low frequency and were not shared across animals. Nineteen unique iSNVs reached majority frequency in at least one sample, (10 nonsynonymous, 8 synonymous, and one variant (PB1 nt 231) that is synonymous in PB1 but nonsynonymous in the overlapping PB1-F2 reading frame). Four of these majority iSNVs were already present in the inoculum at minority frequency (PA L655F, NA G454D, NA S405S, NS1 P164S). Among donor samples, 10 unique iSNVs were detected in multiple donor animals. Five of these shared donor variants were present above the 3% frequency threshold in the inoculum: PA L655F, NA G454D, NA S405S, PB2 S688S, and NS1 P164S. The remaining five shared donor variants were not detected above the 3% threshold in the inoculum and therefore either arose independently in multiple animals or were present in the inoculum below our limit of detection. Only one of these five, M2 I33V, was nonsynonymous. Thus, repeated detection of the same variants across donor animals was driven largely, though not entirely, by standing variation in the inoculum rather than by recurrent within-host emergence. Together, these data indicate that huTX37-H5N1 infections were characterized by limited within-host diversification during acute infection.

We next examined whether HA, a key determinant of mammalian adaptation, exhibited evidence of diversification during infection by analyzing whether nonsynonymous HA mutations were frequent, recurrent, or reached high frequencies. Across both NM93-H5N1– and huTX37-H5N1–infected ferrets, nonsynonymous HA mutations were detected sporadically at low frequency and showed no virus-specific bias (Supplementary Fig. 1 and Supplementary Data 1-2). Only a single HA substitution, I301M (full-length H5 numbering) located in the HA1 region, reached majority frequency by day 3, occurring in one huTX37-H5N1–infected ferret. This mutation has not been functionally characterized and appears to be rare among H5N1 sequences from the U.S. cattle outbreak, although it has been detected sporadically, including in A/cattle/Michigan/24-010071-008/2024.

No nonsynonymous mutations in HA were detected in more than one animal and we observed no consistent enrichment of substitutions previously associated with mammalian receptor adaptation (Fig. 4 and Supplementary Data 1-2). In NM93-H5N1–infected ferrets, one variant of potential interest, HA S136N (corresponding to S120N in mature H5 numbering), was detected at low frequency (∼4%) in a single ferret at 7 days post-infection. Although this substitution has been reported to increase α2,6-linked sialic acid binding in the context of a cold-adapted H5N1 vaccine strain ^27^ and was identified as a minor variant in a pig inoculated with A/Cattle/Texas/063224-24-1/2024 ^28^, deep mutational scanning of H5 HA has not demonstrated enhanced α2,6 binding or viral entry associated with this mutation.^29^ Therefore, its functional significance remains uncertain. We identified no other HA mutations with known or putative functional benefits, including mutations associated with increased α2,6-linked sialic acid binding, in animals infected with either H5N1 virus.

### Genetic drift is detectable from longitudinal iSNV frequencies

The preceding analyses showed that both NM93-H5N1 and huTX37-H5N1 generated limited within-host genetic diversity during ferret infection. We next examined longitudinal iSNV frequency trajectories to determine whether detected variants showed reproducible directional changes across animals or instead displayed allele-frequency fluctuations consistent with genetic drift (Fig. 5 and Supplementary Figures 5–6).

**Figure 5.**
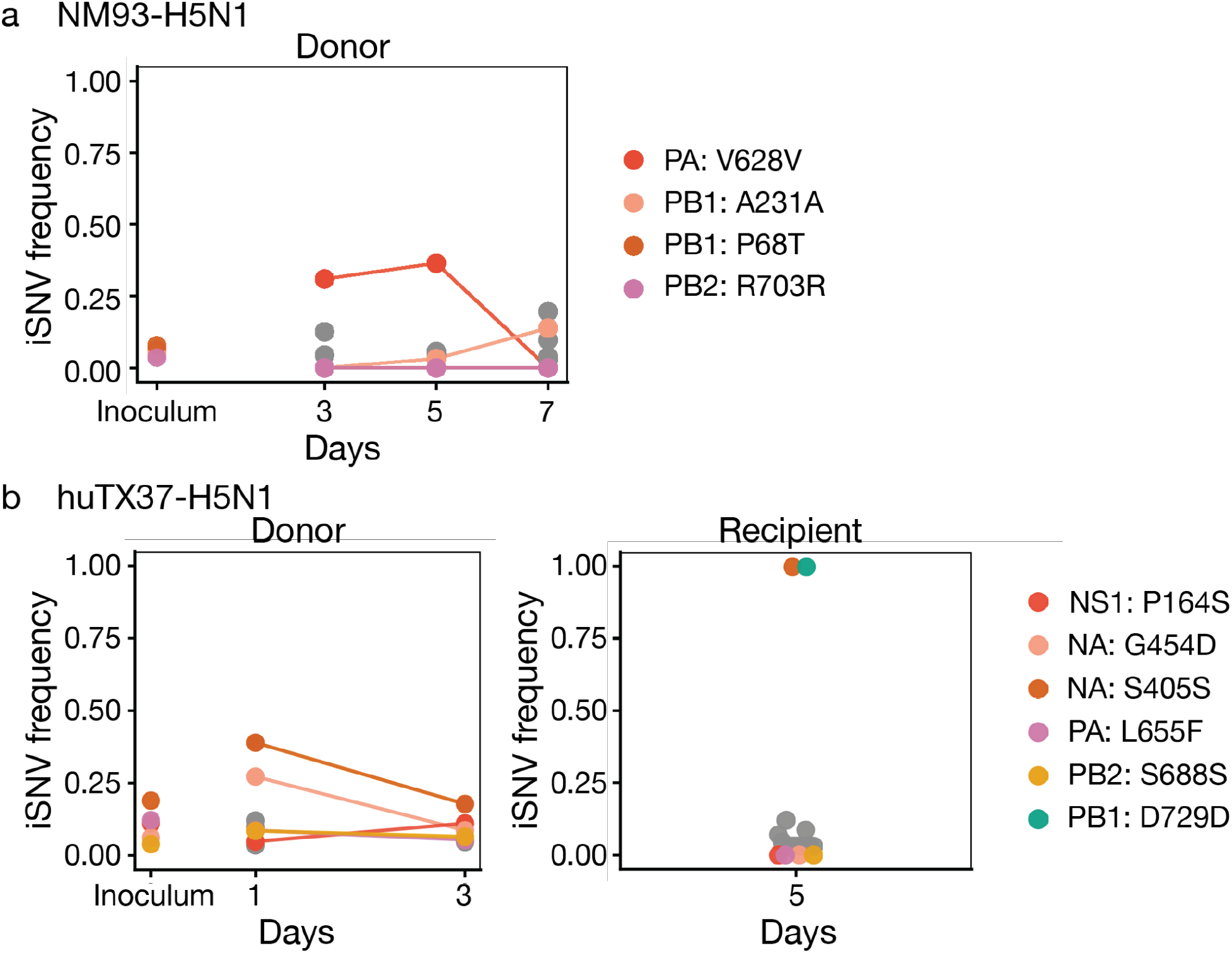
Longitudinal iSNV frequencies across a representative transmission pair for NM93-H5N1 and huTX37-H5N1. (a) NM93-H5N1 iSNV trajectories in stock virus and donor 2.(b) huTX37-H5N1 iSNV trajectories in stock virus, donor and recipient ferret for 1 x10^6^ pair 5. iSNVs found in at least the stock are colored in shades of red and orange; those not present in the stock but present in more than one animal or timepoint, or present at >50% frequency, are in shades of green and blue. Other iSNVs present at a single time point at <50% frequency are displayed in gray.

In the representative NM93-H5N1 donor shown in Fig. 5a, most variants were detected at only a single sampled time point or changed modestly between time points. The inoculum-derived synonymous variant PA V628V increased from 31.0% at day 3 to 36.5% at day 5, but was not detected above the 3% threshold at day 7. A second inoculum-derived variant, PB1 A231A, was detected at 3.2% at day 5 and 13.9% at day 7. Other variants in this animal were detected only transiently. This pattern of intermittent detection and modest, non-monotonic frequency change is consistent with genetic drift, rather than a sustained directional sweep.

The representative huTX37-H5N1 transmission pair shown in Fig. 5b showed similar short-term allele-frequency variability in the donor. In the 1×10⁶ PFU donor from pair 5, all five inoculum-derived variants were recovered at both donor time points, but their frequencies shifted by different amounts between days 1 and 3. NA S405S decreased from 39.0% to 17.8%, NA G454D decreased from 27.2% to 8.7%, and the remaining inoculum-derived variants changed by smaller amounts. In contrast, the recipient sample from the same pair showed a markedly different population composition, with NA S405S near fixation at 99.7%, foreshadowing the strong reshaping of viral diversity during transmission analyzed below.

Across the broader huTX37-H5N1 donor dataset, recovery of inoculum-derived variation depended on inoculation dose. In the 1×10⁶ PFU group, all five inoculum variants were detected in all donor samples, consistent with broad sampling of the stock population during high-dose inoculation. At lower inoculation doses, these same variants were recovered less consistently across donor animals. This dose-dependent heterogeneity is consistent with stochastic sampling of low-frequency inoculum variants during infection, followed by additional allele-frequency fluctuations during within-host replication.

Across the full dataset, large longitudinal frequency changes were uncommon among variants detected above threshold at multiple time points in the same donor. In NM93-H5N1 donors, only 1 of 14 such variant-animal trajectories changed by ≥20 percentage points, while in huTX37-H5N1 donors only 2 of 28 did so. Together, these trajectories provide evidence that genetic drift contributed to within-host allele-frequency changes, while providing little support for repeatable positive selection during acute infection.

### Weak purifying selection is detectable in within-host H5N1 viral populations

To characterize the selective forces shaping viral evolutionary dynamics, we leveraged an existing approach for detecting purifying selection ^30^. In the absence of selection, synonymous and nonsynonymous mutations are expected to accumulate at similar rates, once we adjust for the fact there are more nonsynonymous than synonymous sites. Purifying selection preferentially removes deleterious nonsynonymous mutations. As such, purifying selection can be detected by quantifying the frequencies of detected iSNVs, with nonsynonymous iSNVs expected to be observed at lower frequencies than synonymous iSNVs under purifying selection. To examine evidence for purifying selection, we therefore plotted the proportion of detected iSNVs (across all NM93-H5N1 samples and separately across all huTX37-H5N1 samples) that fall below a given cutoff frequency (Fig. 6). When plotting these proportions, we stratified the iSNVs by whether they were nonsynonymous or synonymous.

**Figure 6.**
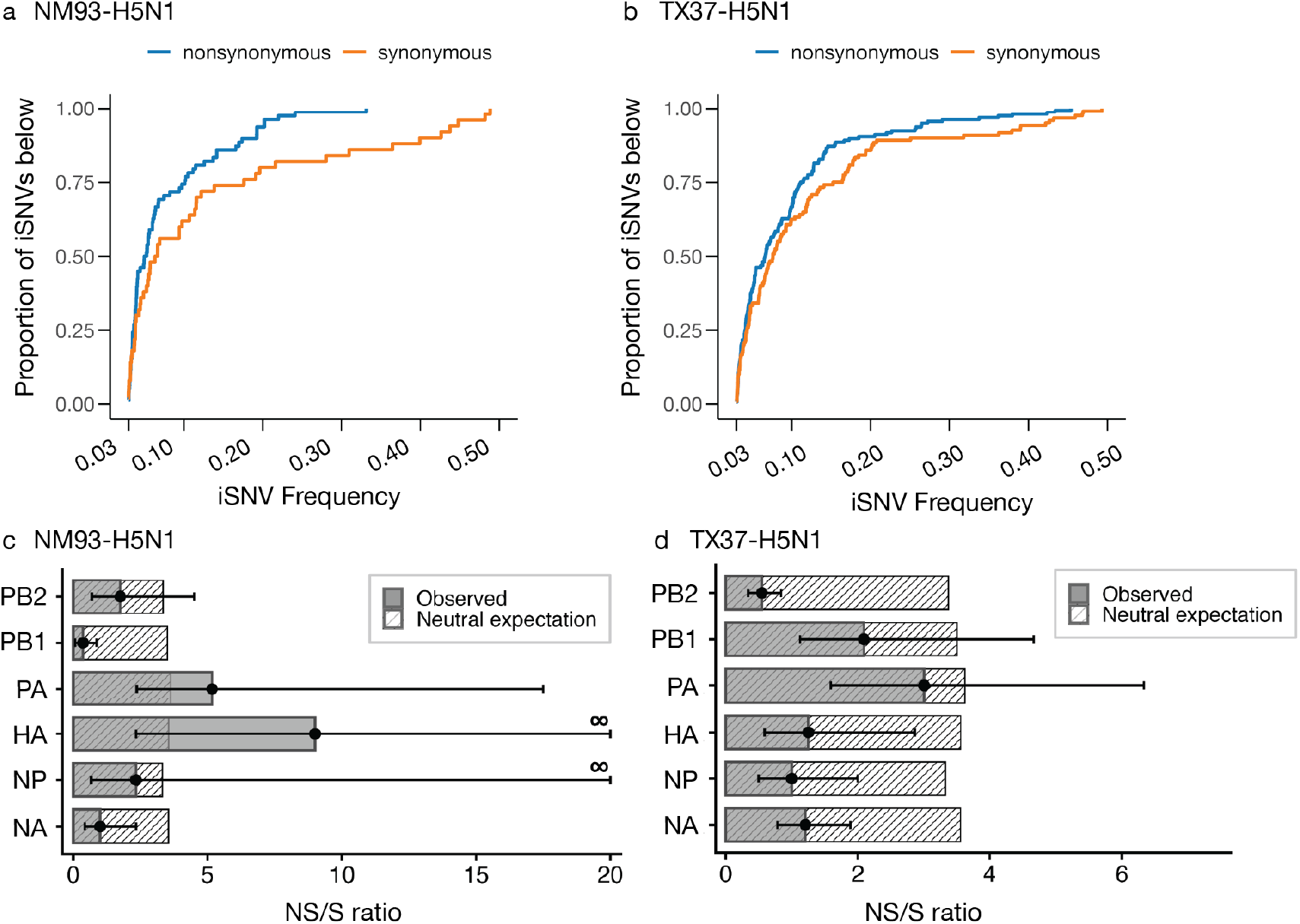
Nonsynonymous iSNV patterns provide evidence of weak within-host purifying selection for NM93-H5N1 and huTX37-H5N1. (a) Cumulative distribution of nonsynonymous and synonymous iSNV frequencies detected in all donor ferrets in the NM93-H5N1 study. (b) Cumulative distribution of nonsynonymous and synonymous iSNV frequencies detected in all ferrets in the huTX37-H5N1 study. (c) The ratio of the number of nonsynonymous to synonymous iSNVs, by gene segment in the NM93-H5N1 study. (d) The ratio of the number of nonsynonymous to synonymous iSNVs, by gene segment, in the huTX37-H5N1 study. For panels (c) and (d), the observed ratios are shown alongside the neutral expectation, given by the ratio of nonsynonymous to synonymous sites. Black whiskers show the 95% confidence interval of the ratio, calculated by 10,000 random draws from a binomial distribution using the observed proportion of nonsynonymous variants.

In animals infected with NM93-H5N1, nonsynonymous iSNVs were disproportionately enriched at very low frequencies relative to synonymous iSNVs, consistent with purifying selection (Fig. 6a). In ferrets infected with huTX37-H5N1, nonsynonymous iSNVs showed only a slight enrichment at lower frequencies compared to synonymous iSNVs, suggesting potentially weaker purifying selection. In addition, we calculated the ratio of non-synonymous to synonymous iSNVs for the major coding gene segments (PB2, PB1, PA, HA, NP, and NA) across all ferrets in the two studies and then compared this ratio to the one expected under neutral evolution ^30^. These N/S ratios showed gene-specific patterns, with confidence intervals for several genes falling significantly below the neutral expectation consistent with purifying selection (Fig. 6c,d). Specifically, PB1 and NA showed evidence of purifying selection in NM93-H5N1, while PB2, HA, NP, and NA showed purifying selection in huTX37-H5N1. The gene-specific evidence of purifying selection in huTX37-H5N1 (Fig. 6d) may help explain why the genome-wide frequency distributions showed only a weak signal of purifying selection (Fig. 6b). Purifying selection may have reduced the number of detectable nonsynonymous iSNVs, leaving a subset of nonsynonymous variants with relatively small fitness costs whose frequencies more closely resembled those of synonymous iSNVs.

As a complementary approach to detect selection, we next quantified nonsynonymous (πN) and synonymous (πS) nucleotide diversity in the 6 major coding genes, which minimizes noise from short and overlapping gene segments, and as we have done previously ^14,15,31^. πN < πS suggests that, on average, purifying selection is acting to remove deleterious mutations from the viral population. Conversely, πN > πS is more consistent with diversifying selection, in which mutations away from the consensus are favored, as might be expected in a virus adapting to a new host. In animals directly inoculated with NM93-H5N1, πN and πS were not significantly different in any genes (Fig. 7a). In donor ferrets infected with huTX37-H5N1, πN was lower than πS in PB2, NP, and NA (Fig. 7b and Supplementary Fig. 2, p = 8.0 × 10⁻^5^, 0.01, and 8.7 × 10⁻^5^, respectively), suggesting that weak purifying selection could have been acting on these genes. Similar patterns of nucleotide diversity were also found in recipient huTX37-H5N1 infected ferrets (Fig. 7c and Supplementary Fig. 2).

**Figure 7.**
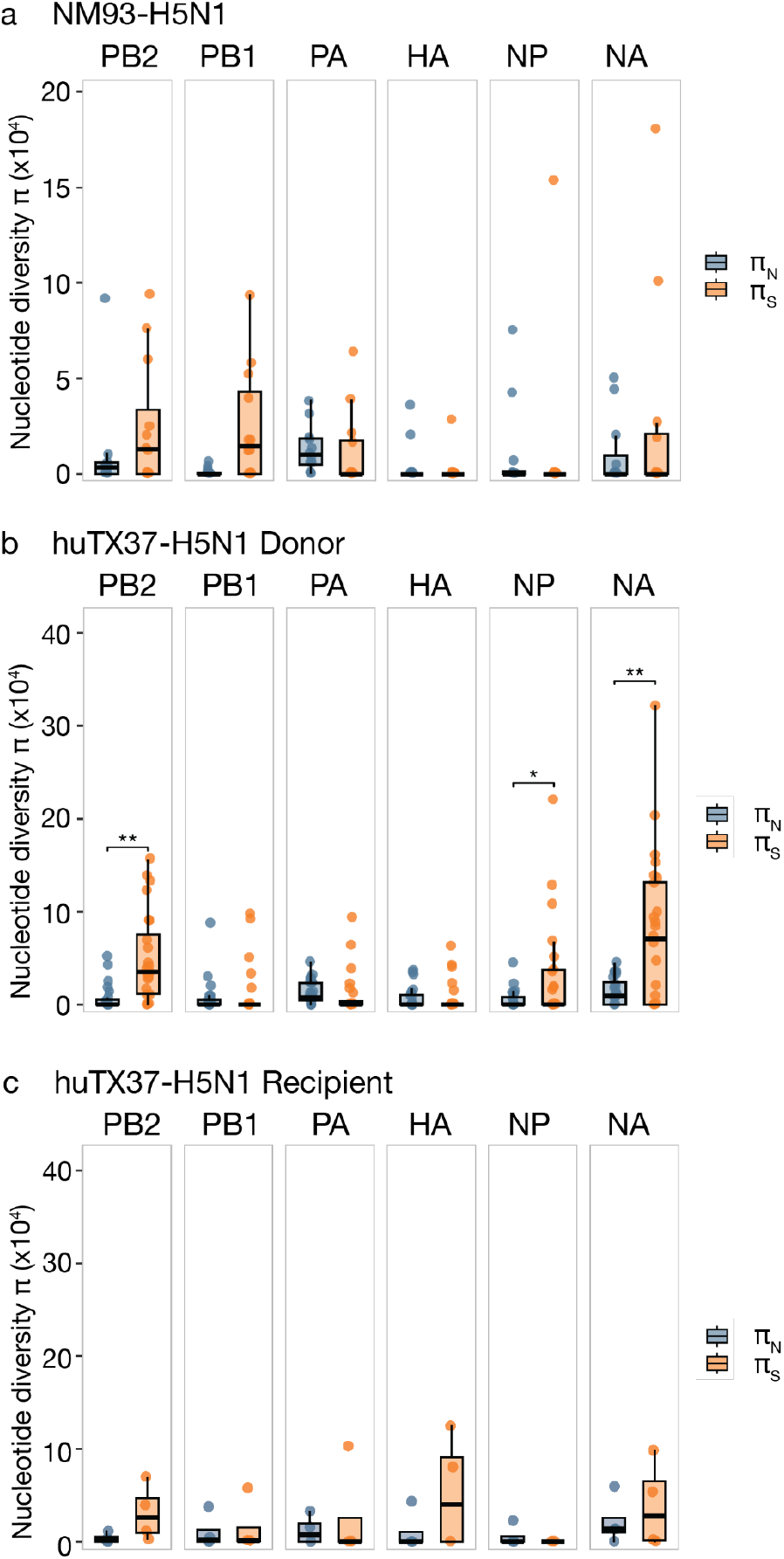
Patterns of within- and between-host evolution assessed using nucleotide diversity (π). (a) πN and πS nucleotide diversity is plotted for the largest ORF on each of the 6 largest gene segments for ferrets infected with NM93-H5N1. (b) πN and πS nucleotide diversity is plotted for each gene segment for ferrets infected with huTx37-H5N1 stratified by donor and (c) recipient ferrets for the earliest available timepoint for each animal. Wilcoxon signed-rank tests were used to compare πN and πS values within each gene segment. Bonferroni correction was applied. Asterisks indicate significance: **\*** p < 0.05; **\*\*** p < 0.01; **\*\*\*** p < 0.001.

Taken together, these within-host summary statistics provide no evidence for diversifying selection and instead support weak purifying selection, superimposed on the stochastic allele-frequency dynamics described above.

### Airborne transmission of huTX37-H5N1 is characterized by a stringent and stochastic transmission bottleneck

The number of viral particles that establish infection in a new host is referred to as the transmission bottleneck size (N_b_). When the bottleneck is tight, a limited subset of viral genetic diversity is transferred from the donor to the recipient. Tight bottlenecks generally act to slow the rate of viral adaptation, as low-frequency variants, including potentially beneficial mutations, may be lost during transmission. In contrast, loose bottlenecks involve a greater number of viruses initiating infection, thereby increasing the likelihood that low-frequency mutations are retained in the new host. Because within-host viral evolution is often dominated by genetic drift and weak purifying selection, beneficial mutations, if they arise, tend to remain at low frequencies within individual hosts, making their transmission particularly sensitive to bottleneck size.

Several statistical methods have been developed to estimate transmission bottleneck size based on the shared genetic variation between donor and recipient pairs ^32–35^. However, these approaches have several important limitations, including an assumption that genetic drift is limited in donors and recipients once viral titers are appreciably high. To avoid needing to make this assumption, we instead applied the bottleneck size estimation approach described by Shi et al. (2024) ^36^which estimates transmission bottleneck size using fixed differences that arise during early viral growth in recipients. In this framework, clonal variants are defined as nucleotide sites where donor and recipient samples are each monomorphic but carry different alleles. The model uses the distribution of clonal variant counts across transmission pairs to jointly estimate the mean transmission bottleneck size and the per-genome mutation rate (μ), conditional on an assumed within-host basic reproduction number (within-host R₀).

To apply this framework to the huTX37-H5N1 experiment, we quantified clonal variants across the five donor-recipient pairs in which airborne transmission was detected, using the same 3% variant-calling threshold applied throughout our analyses. At this threshold, sites with variant frequencies <3% in the donor and >97% in the corresponding recipient were considered clonal variants. We observed three transmission pairs with zero clonal variants, one with a single clonal variant, and one with three clonal variants (Fig. 8D and Supplementary Fig. 3). Because the within-host R₀ has not been quantified in this experimental setting, we estimated the mean N_b_ and the mutation rate μ over a range of plausible within-host R₀ values spanning from 1.5 to 20 ^37^.

**Figure 8.**
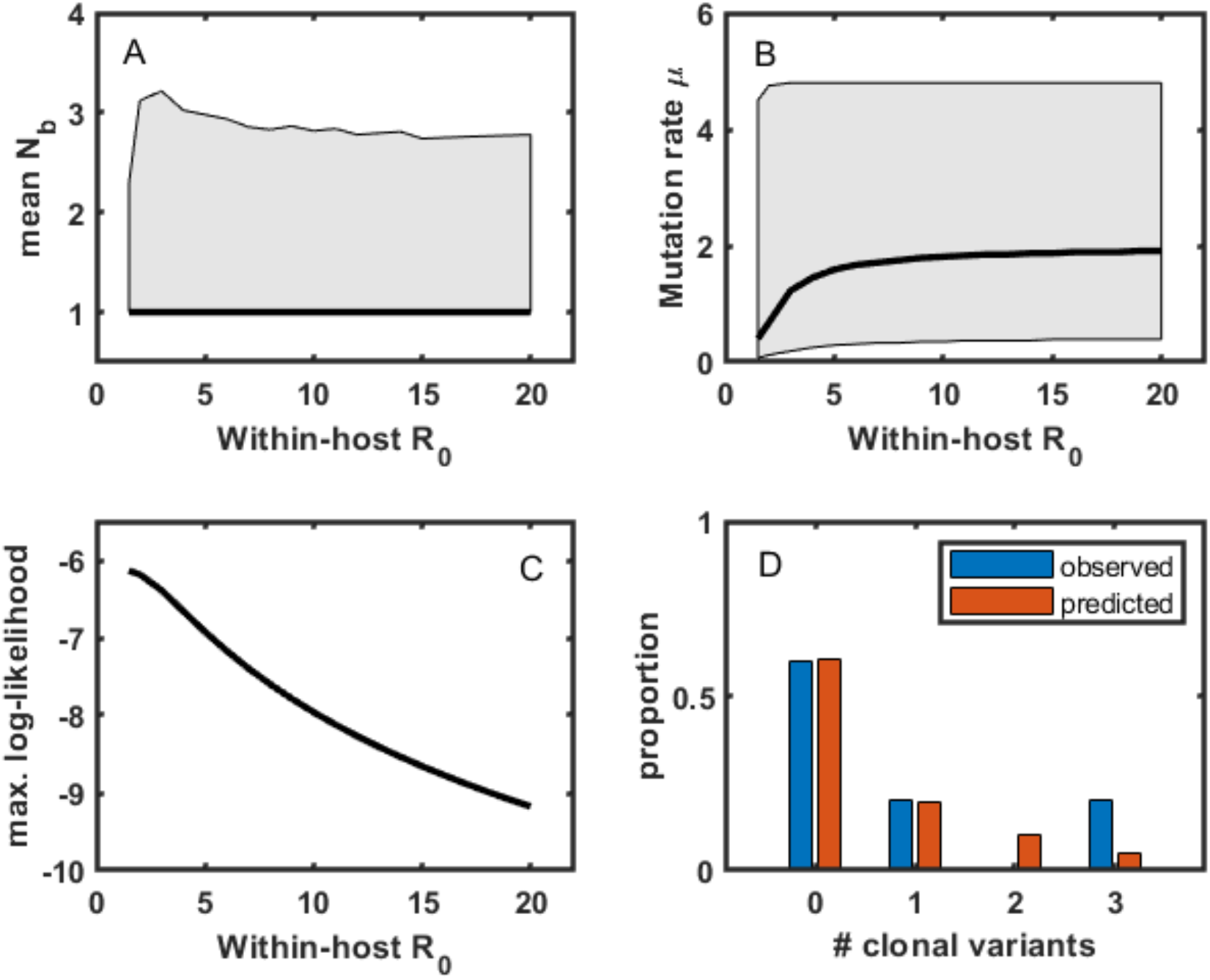
Transmission bottleneck N_b_ estimation for huTX37-H5N1 using clonal variants infers a stringent bottleneck. (A) Estimates of mean transmission bottleneck sizes across a range of plausible within-host basic reproduction numbers. (B) Estimates of the per genome, per infection cycle mutation rate across the same range of plausible within-host basic reproduction numbers. (C) Log-likelihood values of model fits, with models parameterized according to the within-host basic reproduction numbers on the x-axis and with corresponding mean N_b_ and μ values from panels (A) and (B). (D) Estimated proportion of transmission pairs by their number of clonal variants plotted alongside observed proportions. Proportions were estimated using model parameters that yielded the highest likelihood: within-host R₀ = 1.50, mean N_b_ = 1.00, and μ = 0.40 per genome per infection cycle.

Across this range of within-host R₀ values, the mean N_b_ estimate was close to 1 across the entire range of within-host R₀ values considered (Fig. 8a). These estimates are consistent with prior evidence that influenza A virus transmission often involves narrow bottlenecks, including estimates from natural human influenza A transmission and experimental ferret studies showing stringent bottlenecks during mammalian transmission of avian influenza viruses ^14,15,35,36^. Mutation rate estimates were sensitive to the within-host R₀ assumed, ranging from a minimum of 0.40 mutations per genome per infection cycle at an R₀ of 1.5 to a maximum of 1.91 mutations per genome per infection cycle at an R₀ of 20 (Fig. 8b). On a per site level, these estimates are also broadly consistent with mutation rates estimated for human influenza A virus ^38^. Model likelihoods were higher at lower values of within-host R₀, with a within-host R₀ of 1.50 yielding the highest log-likelihood (Fig. 8c). This low best-fit within-host R₀ could reflect a truly low within-host R₀ for huTX37-H5N1, or it could reflect model sensitivity to extensive intercellular heterogeneity in viral progeny production within infected hosts, which could affect the accumulation of clonal variants. We next compared the distribution of clonal variants predicted by this best-fit model to the observed one to assess whether the model was able to reproduce the distribution of clonal variants observed in the data. The model-predicted distribution of clonal variants closely recapitulated the observed data (Fig. 8d, red bars).

We next asked whether transmission was associated with consistent selective filtering of viral genetic diversity. Across the five transmission pairs, πN and πS did not differ significantly between donors and recipients for any gene segment at the earliest available time points, and similar patterns were observed when all time points were considered (Fig. 7b,c and Supplementary Fig. 2). These data provide no evidence for gene-specific selective filtering during transmission. This interpretation is also consistent with the observed iSNV trajectories: most donor variants were not recovered in recipient ferrets, and the few variants detected in recipients included both synonymous and nonsynonymous mutations as well as both inoculum-derived and newly detected variants.

Two inoculum-derived variants, NA S405S and NA G454D, were detected in more than one recipient, although only NA S405S was near-fixed in multiple recipients. Eight recipient variants reached majority frequency, six of which exceeded 97% frequency. No newly detected fixed mutation was shared across recipients, and no recipient variant has previously been associated with mammalian adaptation. Thus, airborne transmission of huTX37-H5N1 appeared highly stringent and largely stochastic, reflecting the sampling of a small number of founding genomes rather than consistent selection for specific adaptive variants.

## Discussion

Wild birds are the reservoir for almost all influenza A viral genetic diversity, and avian influenza viruses pose epizootic and pandemic threats. The evolutionary pathways by which avian influenza viruses adapt to enable efficient infection and/or transmission in mammals remain poorly understood. The 2024-2025 outbreak in US dairy cattle further underscores that avian influenza viruses can emerge unpredictably, even in hosts not previously known to support influenza A virus replication. In this study, we examined the within- and between-host evolutionary dynamics of two bovine-derived clade 2.3.4.4b B3.13 genotype H5N1 viruses, NM93-H5N1 and huTX37-H5N1, whose replication, transmission, and pathogenic potential in ferrets were previously characterized. Like other bovine-derived H5N1 viruses, these isolates harbored mammal-adapting mutations in their polymerase genes but retain HA proteins with primarily avian-like receptor-binding phenotypes. Here we asked whether there was any evidence that natural selection might favor additional mammal adaptation as these viruses replicated and transmitted in ferrets. Within-host viral genetic diversity was limited in all animals with detectable infectious virus, and viral evolution was shaped by genetic drift and weak purifying selection. We found no evidence of additional mammalian adaptation in either experiment. Productive airborne transmission, when detected for huTX37-H5N1, involved very narrow bottlenecks, with no evidence for natural selection on particular gene segments. Moreover, our evolutionary analyses did not reveal a clear genetic or within-host evolutionary explanation for why airborne exposure resulted in detectable productive infection in some recipient animals but not others. Together, our results suggest that stochastic processes acting within hosts and during transmission can constrain the acquisition and propagation of additional mammalian-adaptive changes, including HA substitutions that could improve α2,6-linked receptor binding and enhance airborne transmissibility.

The upper respiratory tracts of humans and ferrets are enriched for α2,6-linked sialic acids, whereas avian influenza viruses preferentially bind α2,3-linked sialic acids. Accordingly, a shift toward α2,6-linked receptor binding has been shown to be a key determinant of efficient airborne transmission in mammals ^11^. Bovine-derived H5N1 viruses provide a particularly relevant context in which to examine this evolutionary pressure. Although both α2,3- and α2,6-linked sialic acids are present in bovine mammary tissues ^39^, NM93-H5N1 and huTX37-H5N1 retain primarily avian-like receptor-binding phenotypes, indicating that efficient replication in cattle-associated tissues does not necessarily require a shift toward dominant α2,6-linked receptor binding. In contrast, the ferret upper respiratory tract contains abundant α2,6-linked receptors and provides an environment in which HA mutations that enhance mammalian-type receptor binding could plausibly be favored, even during replication in directly inoculated animals. Prior work from our group showed that avian-like influenza viruses can acquire or enrich HA mutations during ferret infection and transmission, including variants present below 10% frequency in donors ^14,15^. We therefore might have expected these viruses to acquire or enrich HA mutations associated with enhanced mammalian receptor binding during replication and transmission in ferrets. In contrast, we observed no signs of such adaptation: no recurrent HA mutations, no enrichment of nonsynonymous variation in HA, no evidence of HA-focused diversification during infection, and no selective sweep in HA during transmission.

The absence of detectable HA adaptation can be interpreted through two different types of evolutionary constraints: mutation limitation and selection limitation. Under a mutation-limited scenario, receptor-adapting mutations may simply fail to arise during short acute infections, particularly if the mutational target in HA is small or if receptor-switching phenotypes require combinations of mutations rather than a single substitution. Although individual HA mutations can increase binding to α2,6-linked sialic acids in some genetic backgrounds, such mutations may impose fitness costs by decreasing HA stability, altering glycosylation, or disrupting HA/NA balance ^27,40,41^. In this case, receptor-adapting variants would require additional compensatory changes to preserve replication fitness, reducing the probability that a fit, transmissible genotype emerges during a single acute infection. Together, the limited standing diversity in both inocula and the limited *de novo* diversification observed during infection support a role for mutation limitation: the viruses began infection with little starting genetic material for within-host diversification or immediate selection, and few additional candidate adaptive mutations were generated during the sampled acute infection window.

Under a selection-limited scenario, mammal-adapting variants may arise but fail to increase to detectable frequencies because the selective advantage they confer is weak, spatially restricted, or counteracted by stochastic processes. Several observations are consistent with this possibility. First, both NM93-H5N1 and huTX37-H5N1 replicated to high titers in directly inoculated ferrets, indicating that these viruses are capable of robust replication in the ferret respiratory tract. Second, ferret airways contain α2,3-linked sialic acids on multiple epithelial cell types ^42,43^, and bovine-derived H5 HAs may retain limited capacity to utilize α2,6-linked receptors ^10,16^. Thus, the founding avian-like viral population may already exploit the available within-host environment sufficiently well that receptor-adapting variants have only a modest advantage, if any, during acute infection. Limited airborne transmission may therefore be possible even without complete receptor switching, while inefficient mammalian-type receptor binding may still remain an important barrier to sustained transmission and pandemic emergence.

Spatial structure within infected hosts could further limit the efficiency of selection. Direct intranasal inoculation, particularly at high dose, may seed many susceptible sites with the founding genotype early in infection. If an α2,6-adapted variant arises later, it may do so within a local target-cell population that is already dominated by the ancestral genotype or in an anatomical region where its phenotypic advantage is limited. This idea is analogous to ecological priority effects, in which early-established populations can occupy available niches and limit the expansion of later-arriving competitors unless those competitors have a large immediate advantage ^44–46^. Here, the analogy applies within individual donor animals rather than among competing donors or at the level of recipient colonization. Prior work using barcoded influenza viruses showed that replication can occur in spatially restricted “islands” of infection, with genetically distinct viral populations in different lung lobes ^47^. Such compartmentalization means the respiratory tract is not a single well-mixed pool of virus; a beneficial variant that arises in one anatomical region may not efficiently access the upper-airway niche where it would be most strongly favored for onward transmission. Spatial structure could therefore dull selection by limiting mixing among viral subpopulations and by separating the origin of a beneficial mutation from the site where its advantage would be most relevant.

These findings place the idea that mammalian replication and airborne transmission are related but partially separable phenotypes in an evolutionary context. Both viruses examined here achieved high viral loads in directly inoculated animals, yet outcomes in exposed recipients were limited: NM93-H5N1 produced serologic evidence of exposure in one recipient without detectable infectious virus, whereas huTX37-H5N1 produced detectable productive infection in a subset of exposed recipients. For both isolates, within-host evolution was characterized by limited diversification, genetic drift, and weak purifying selection. Thus, efficient replication in mammals did not coincide with efficient productive airborne transmission or detectable evolutionary movement toward enhanced airborne spread. This disconnect may reflect both mutation limitation, in which few fit transmission-enhancing variants are generated, and selection limitation, in which the acute within-host environment does not impose strong selection for new phenotypes such as enhanced α2,6-linked receptor binding.

Stringent transmission bottlenecks provide an additional constraint on onward evolution of bovine-derived H5N1 in mammals. We estimated mean bottleneck sizes close to one virion, indicating that most within-host diversity is lost during airborne transmission. This sharply limits propagation of rare variants, including those with modest beneficial effects, between hosts. In principle, narrow bottlenecks can reshuffle allele frequencies and occasionally reset the competitive context by allowing a rare genotype to found infection in a new host with reduced competition from the ancestral genotype ^34^. Prior ferret transmission studies from our group showed that transmission can impose strong selection on HA and rapidly alter viral population composition ^14,15^. In the present study, however, we found no analogous evidence of selection on any gene segment: there was no recurrent fixation of a newly detected adaptive variant, no gene-specific reduction in diversity consistent with selective filtering, and no enrichment of mutations associated with mammalian adaptation in recipients. One explanation is that donor populations contained little phenotypic diversity for transmission selection to act on. Thus, transmission appeared to act primarily as a stochastic filter in this system, while still leaving open the possibility that rare selective transmission events could occur under different conditions or with a larger number of transmission events.

Several limitations should be considered when interpreting these results. The number of successful transmission events was small, limiting statistical power to detect rare or weakly selected variants. If enhanced transmission fitness arises rarely and unpredictably, it is unsurprising that we observed no such events among five transmissions. In addition, these were acute infections, and huTX37-H5N1 infections were rapidly fatal, shortening the period during which viral diversification, within-host selection, and transmission could occur.

Together our findings suggest that additional mammal-adapting changes may be unlikely to arise predictably within any single acute mammalian infection. Evolutionary trajectories toward efficient mammalian transmission appear constrained. Possible factors driving these constraints include genetic context and functional tradeoffs that limit the emergence of fit adaptive variants ^27,40,41^; spatial structure and stochastic dynamics that reduce the efficiency of within-host selection ^31,47^; and stringent between-host bottlenecks that limit onward propagation of rare variants ^14,15,35,36^. However, such adaptation remains possible and may not require canonical markers of mammalian adaptation. The occurrence of multiple H5N1 introductions into cattle demonstrates the ongoing potential for agricultural spillover events. The bovine outbreak experience underscores that influenza A viruses have broad host tropism, and viruses with clade 2.3.4.4b HA remain a pandemic and epizootic threat. Continued circulation of viruses from this clade creates repeated opportunities for rare but consequential evolutionary jackpot events to emerge and be amplified. Continued assessment of zoonotic risk therefore remains essential, and sustained genomic surveillance of clade 2.3.4.4b influenza viruses should be considered a priority.

## Methods

### Study description and samples analyzed

To investigate within-host and between-host viral evolutionary dynamics during mammalian transmission of bovine-derived H5N1 influenza A viruses, we analyzed samples from two independent ferret transmission experiments. The first study evaluated transmission of bovine H5N1 virus A/dairy cattle/New Mexico/A240920343-93/2024 (NM93-H5N1) in ferrets ^10^ but did not include viral sequencing data; nasal swab samples and inoculum from that study were therefore sequenced as part of the present work. The second study evaluated transmission of bovine H5N1 virus A/Texas/37/2024 (huTX37-H5N1) in ferrets and reported deep sequencing of viral inoculum stocks and longitudinal nasal swab samples; the publicly available sequencing data generated in that study were reanalyzed here using a uniform bioinformatic pipeline to enable direct comparison ^9^.

Across both studies, viral RNA from inoculum stocks and from nasal swabs collected longitudinally from experimentally infected (donor) and recipient ferrets was analyzed. Only samples with viral loads above the assay limit of detection were included for sequencing and downstream evolutionary analyses. Sample identifiers, associated metadata, and corresponding SRA accession numbers for all sequences analyzed in this study are provided in Supplemental Table 1.

All samples used for sequencing analysis were inactivated in biosafety level 3 containment laboratories at the Influenza Research Institute of the University of Wisconsin-Madison prior to sequencing at biosafety level 2. All experiments were approved by Institutional Biosafety Committees of the University of Wisconsin-Madison. Funding for this study came in part from the NIAID Centers of Excellence for Influenza Research and Response (contract no. 75N93021C00014) that supported the previously published research ^9, 10^.The NIAID grant was reviewed by the University of Wisconsin-Madison Dual Use Research of Concern (DURC) Subcommittee in accordance with the May 2024 United States Government Policy for Oversight of Dual Use Research of Concern and Pathogens with Enhanced Pandemic Potential and determined to not meet the criteria of DURC or PEPP. The University of Wisconsin-Madison Institutional Contact for Dual Use Research reviewed this manuscript and confirmed that the studies described herein do not meet the criteria of DURC or PEPP.

### Viral genome sequencing

Total RNA was extracted from aliquots of inoculum used to infect ferrets, or from ferret nasal swab samples with either the QIAamp Viral RNA Mini kit (Qiagen) or MagMAX-96 viral RNA isolation kit (Invitrogen) according to the respective manufacturer’s instructions. Single-reaction amplification of viral genomes was performed using pooled gene-specific primers (Supplementary Table 1) as previously described ^9^. Amplicon libraries were fragmented using an Illumina DNA Prep Kit (Illumina) and indexed with Illumina DNA/RNA UD Indexes (Illumina), according to the manufacturer’s instructions. DNA libraries were sequenced on an Illumina MiSeq system using MiSeq Reagent Kit v.3 (600 cycle) cartridges (Illumina). All samples sequenced as part of the present study were processed using the same library preparation and sequencing protocols to minimize technical variability.

### Processing raw sequence data, mapping, variant calling and variant annotation

A bespoke Nextflow pipeline was used to quality filter reads, align reads, and call variants (https://github.com/nrminor/oneroof). The pipeline reproducibly supplies software dependencies via Docker or Apptainer containers, or via local conda environments built with the Pixi package manager. For the results presented here, we required a minimum read length of 150 bases and a maximum read length of 450 bases, along with a minimum average read quality of 20. Reads were aligned to isolate-matched reference genomes to minimize reference bias using minimap2 v2.28. Reference genomes were A/dairy cattle/New Mexico/A240920343-93/2024 (GenBank accessions: PQ067970, PQ068570, PQ068393, PQ068523, PQ068538, PQ068546, PQ068553, and PQ068562) and A/Texas/37/2024 (GenBank accessions: PP577947, PP577941, PP577942, PP577943, PP577945, PP577944, PP577946, and PP577940). iVar v1.4.2 ^48^ was used to call variants and snpEff v5.2 was used for variant-effect annotation. Variants were called using a minimum required coverage depth of 200 reads and a minimum variant frequency of 0.03. This conservative frequency threshold was chosen to reduce the contribution of sequencing error and falls within the range used in prior intrahost respiratory virus evolution studies ^23,24^. Variants at a frequency of ≥0.5 were considered majority-level (consensus), and those at 0.03–0.5 were considered minor (sub-consensus) intrahost single-nucleotide variants (iSNVs).

### Consensus sequence generation

Initial trimming and filtering of Illumina MiSeq reads were performed using Local Run Manager v.3.0.0. Demultiplexed reads were processed using the iterative refinement meta-assembler IRMA v.1.0.2 to generate consensus sequences for each influenza virus gene segment. Default IRMA FLU parameters were used, except that LABEL was used for read sorting, the residual assembly factor was set to 400 for the secondary assembly, and reference elongation was prevented. Consensus sequences were obtained from the secondary assembly.

### Cumulative proportion iSNV plots

We generated cumulative proportion plots of synonymous and nonsynonymous intrahost single-nucleotide variants (iSNVs) across the viral genome, as previously described ^30^. For each virus dataset (NM93-H5N1 and huTX37-H5N1), all iSNVs with allele frequencies below 50% were pooled across samples and classified as synonymous or nonsynonymous based on snpEff annotation. Analyses included variants annotated to the following coding regions: PB2, PB1, PB1-F2, PA, PA-X, HA, NP, NA, M1, M2, NS1, and NS2. These plots were intended as a descriptive, genome-wide summary; therefore all annotated coding regions were included, including overlapping segments.

For each variant class, the empirical cumulative distribution function (ECDF) of iSNV frequencies was calculated, such that the y-axis represents the cumulative proportion of iSNVs with frequencies less than or equal to a given value on the x-axis. Because variant effects were annotated independently for each coding region, mutations occurring in overlapping reading frames (e.g., PA and PA-X; PB1 and PB1-F2; M1 and M2; NS1 and NS2) contribute to each relevant coding annotation and are therefore counted multiple times in this genome-wide analysis.

### Ratio of nonsynonymous to synonymous iSNVs

To quantify the relative abundance of nonsynonymous and synonymous variation, we calculated the ratio of the number of observed nonsynonymous iSNVs to the number of observed synonymous iSNVs (N/S ratio) on a per-gene basis. For each gene (PB2, PB1, PB1-F2, PA, PA-X, HA, NP, and NA), iSNVs were pooled across all ferrets separately for NM93-H5N1 and huTX37-H5N1 and classified based on snpEff annotation. Variants occurring in overlapping reading frames were counted independently for each affected gene.

To assess whether observed N/S ratios deviated from neutral expectations, we compared them to expected ratios based on codon-level site availability. Expected N/S ratios were calculated using the ratio of nonsynonymous to synonymous sites (N_sites/S_sites) for each gene, as estimated by SNPGenie. Ninety-five percent confidence intervals for observed N/S ratios were generated using 10,000 binomial resamples parameterized by the observed proportion of nonsynonymous iSNVs.

The M and NS segments were excluded from this analysis because of their short length, extensive overlapping coding regions, and limited numbers of independently callable sites which reduce statistical power and complicate interpretation of N/S ratios for these segments.

### Nucleotide diversity calculations

Per-gene nucleotide diversity was quantified using π summary statistics. Nucleotide diversity (π) measures the average number of pairwise nucleotide differences per site within a population and was calculated using SNPGenie (https://github.com/chasewnelson/SNPGenie) ^49^. SNPGenie uses a codon-based framework derived from the Nei and Gojobori method to estimate overall nucleotide diversity (π) and to partition diversity into synonymous (πS) and nonsynonymous (πN) components from next-generation sequencing variant data.

SNPGenie was provided with reference genome sequences, gene annotations, and variant calls generated as described above. Positions annotated as mixed or ambiguous in the reference sequence were excluded from analysis. Nucleotide diversity was calculated for PB2, PB1, PA, HA, NP, and NA.

The M and NS segments were excluded from nucleotide diversity analyses because of their short length, extensive overlapping coding regions, and limited numbers of callable sites, which reduce statistical power and complicate interpretation of π for these segments.

### Statistical comparisons

Statistical comparisons were performed in R using the base-R wilcox.test function. For non-normally distributed data, we employed non-parametric statistical tests. We used paired Wilcoxon signed-rank tests for matched (dependent) samples and unpaired Wilcoxon rank-sum tests for unmatched samples.

### Transmission bottleneck estimation using clonal variants

Mean transmission bottleneck sizes (*N_b_*) were estimated using the clonal variant method of Shi et al. (2024)^36^. Briefly, we identified fixed *de novo* mutations in each recipient and used their frequency distribution across transmission pairs to infer mean *N_b_* under a stochastic branching-process model of early within-host viral growth.

For each donor–recipient pair, clonal variants were identified as those that are monomorphic in both the donor and recipient but carry different alleles (using 3% variant frequency thresholds). The observed distribution of clonal variants was then fitted via maximum likelihood to a model that jointly estimates the mutation rate (μ) and mean bottleneck size (*N_b_*), assuming early exponential viral expansion with within-host reproduction number R₀.

### Data visualization and figure preparation

All statistical analyses and data visualizations were performed in R (version 4.4.2) and Matlab. Schematic figures were created using BioRender.

## Data availability

Deep-sequencing data have been deposited to the Sequence Read Archive (SRA) under bioproject accession no. PRJNA1163435 for huTX37-H5N1. Deep-sequencing data for NM93-H5N1 are in process of being uploaded to SRA. SRA accession numbers for individual samples are provided in Supplementary Data 3. Consensus sequences for NM93-H5N1 have been deposited in Genbank and accession numbers are available in Supplementary Data 4. Transmission bottleneck analysis code is available at https://github.com/koellelab/H5N1_ferrets/ and remaining analysis code is available at https://github.com/hmmachko/bovine_H5N1_ferret_transmission/tree/main/.

## Author contributions

WW contributed to data curation, methodology, data analysis, data visualization, and writing of the initial draft. HM contributed to methodology, data analysis, data visualization, writing, draft review, and editing. TCF contributed to conceptualization, methodology, data analysis, writing-draft review and editing, and supervision. KK contributed to data analysis and writing-draft review and editing. CG, LG, AB, TM, TW, LB, PJH, AE, GN, and YK reviewed and edited the manuscript.

## Acknowledgements

The authors would like to thank Nick Minor for the generation of the preprocessing and alignment of influenza sequencing data, which was utilized for this study.

H.M.M. received funding from US National Institutes of Health (grant number T32AI055397) and the Pearl Stetler Fellowship. This work was supported by the National Institute of Allergy and Infectious Diseases Centers of Excellence for Influenza Research and Response (contract 75N93021C00014).

## Supplementary Information

Supplementary Information includes Supplementary Figs. 1–6, Supplementary Table 1, and descriptions of Supplementary Data 1–4. Supplementary Data 1–4 are provided as separate tabular CSV files.

## References

1. HPAI Confirmed Cases in Livestock. Animal and Plant Health Inspection Service https://www.aphis.usda.gov/livestock-poultry-disease/avian/avian-influenza/hpai-detections/hpai-confirmed-cases-livestock.

2. Update: Genetic Sequencing Results for Wisconsin Dairy Herd Detection of Highly Pathogenic Avian Influenza. Animal and Plant Health Inspection Service https://www.aphis.usda.gov/news/agency-announcements/update-genetic-sequencing-results-wisconsin-dairy-herd-detection-highly (2025).

3. Caserta, L. C. et al. Spillover of highly pathogenic avian influenza H5N1 virus to dairy cattle. Nature 634, 669–676 (2024).

4. Guan, L. et al. Cow’s milk containing avian influenza A(H5N1) virus - heat inactivation and infectivity in mice. N. Engl. J. Med. 391, 87–90 (2024).

5. CDC. H5 Bird Flu: Current Situation. Avian Influenza (Bird Flu) https://www.cdc.gov/bird-flu/situation-summary/index.html (2025).

6. Garg, S. et al. Outbreak of highly pathogenic avian influenza A(H5N1) viruses in U.s. dairy cattle and detection of two human cases - United States, 2024. MMWR Morb. Mortal. Wkly. Rep. 73, 501–505 (2024).

7. DSHS reports first human case of avian influenza in Texas. https://www.dshs.texas.gov/news-alerts/dshs-reports-first-human-case-avian-influenza-texas.

8. Pulit-Penaloza, J. A. et al. Transmission of a human isolate of clade 2.3.4.4b A(H5N1) virus in ferrets. Nature 636, 705–710 (2024).

9. Gu, C. et al. A human isolate of bovine H5N1 is transmissible and lethal in animal models. Nature 636, 711–718 (2024).

10. Eisfeld, A. J. et al. Pathogenicity and transmissibility of bovine H5N1 influenza virus. Nature 633, 426–432 (2024).

11. Matrosovich, M. et al. Early alterations of the receptor-binding properties of H1, H2, and H3 avian influenza virus hemagglutinins after their introduction into mammals. J. Virol. 74, 8502–8512 (2000).

12. de Graaf, M. & Fouchier, R. A. M. Role of receptor binding specificity in influenza A virus transmission and pathogenesis. EMBO J. 33, 823–841 (2014).

13. Lin, T.-H. et al. A single mutation in bovine influenza H5N1 hemagglutinin switches specificity to human receptors. Science 386, 1128–1134 (2024).

14. Moncla, L. H. et al. Selective bottlenecks shape evolutionary pathways taken during mammalian adaptation of a 1918-like avian influenza virus. Cell Host Microbe 19, 169–180 (2016).

15. Wilker, P. R. et al. Selection on haemagglutinin imposes a bottleneck during mammalian transmission of reassortant H5N1 influenza viruses. Nat. Commun. 4, 2636 (2013).

16. Song, H. et al. Receptor binding, structure, and tissue tropism of cattle-infecting H5N1 avian influenza virus hemagglutinin. Cell 188, 919–929.e9 (2025).

17. Zhang, X. et al. Enhanced pathogenicity and neurotropism of mouse-adapted H10N7 influenza virus are mediated by novel PB2 and NA mutations. J. Gen. Virol. 98, 1185–1195 (2017).

18. Patrono, L. V. et al. Archival influenza virus genomes from Europe reveal genomic variability during the 1918 pandemic. Nat. Commun. 13, 2314 (2022).

19. Dholakia, V. et al. Polymerase mutations underlie early adaptation of H5N1 influenza virus to dairy cattle and other mammals. Microbiology (2025).

20. Mostafa, A. et al. Avian influenza A (H5N1) virus in dairy cattle: origin, evolution, and cross-species transmission. MBio 15, e0254224 (2024).

21. Subbarao, E. K., London, W. & Murphy, B. R. A single amino acid in the PB2 gene of influenza A virus is a determinant of host range. J. Virol. 67, 1761–1764 (1993).

22. Hatta, M., Gao, P., Halfmann, P. & Kawaoka, Y. Molecular basis for high virulence of Hong Kong H5N1 influenza A viruses. Science 293, 1840–1842 (2001).

23. Braun, K. M. et al. Acute SARS-CoV-2 infections harbor limited within-host diversity and transmit via tight transmission bottlenecks. PLoS Pathog. 17, e1009849 (2021).

24. Grubaugh, N. D. et al. An amplicon-based sequencing framework for accurately measuring intrahost virus diversity using PrimalSeq and iVar. Genome Biol 20, 8 (2019).

25. Kim, J. H. et al. Role of host-specific amino acids in the pathogenicity of avian H5N1 influenza viruses in mice. J. Gen. Virol. 91, 1284–1289 (2010).

26. Omoto, S. et al. Characterization of influenza virus variants induced by treatment with the endonuclease inhibitor baloxavir marboxil. Sci. Rep. 8, 9633 (2018).

27. Wang, W. et al. Glycosylation at 158N of the hemagglutinin protein and receptor binding specificity synergistically affect the antigenicity and immunogenicity of a live attenuated H5N1 A/Vietnam/1203/2004 vaccine virus in ferrets. J. Virol. 84, 6570–6577 (2010).

28. Kwon, T. et al. Pathogenicity and transmissibility of bovine-derived HPAI H5N1 B3.13 virus in pigs. bioRxiv 2025.03.04.641414 (2025) doi:10.1101/2025.03.04.641414.

29. Dadonaite, B. et al. Deep mutational scanning of H5 hemagglutinin to inform influenza virus surveillance. PLoS Biol. 22, e3002916 (2024).

30. VanInsberghe, D. et al. Genetic drift and purifying selection shape within-host influenza A virus populations during natural swine infections. PLoS Pathog. 20, e1012131 (2024).

31. Braun, K. M. et al. Avian H7N9 influenza viruses are evolutionarily constrained by stochastic processes during replication and transmission in mammals. Virus Evol. 9, vead004 (2023).

32. Ghafari, M., Lumby, C. K., Weissman, D. B. & Illingworth, C. J. R. Inferring transmission bottleneck size from viral sequence data using a novel haplotype reconstruction method. J. Virol. 94, (2020).

33. Emmett, K. J., Lee, A., Khiabanian, H. & Rabadan, R. High-resolution genomic surveillance of 2014 ebolavirus using shared subclonal variants. PLoS Curr. 7, (2015).

34. Zwart, M. P. & Elena, S. F. Matters of size: Genetic bottlenecks in virus infection and their potential impact on evolution. Annu. Rev. Virol. 2, 161–179 (2015).

35. Sobel Leonard, A., Weissman, D. B., Greenbaum, B., Ghedin, E. & Koelle, K. Transmission bottleneck size estimation from pathogen deep-sequencing data, with an application to human influenza A virus. J. Virol. 91, (2017).

36. Shi, Y. T., Harris, J. D., Martin, M. A. & Koelle, K. Transmission bottleneck size estimation from DE Novo viral genetic variation. Mol. Biol. Evol. 41, msad286 (2024).

37. Baccam, P., Beauchemin, C., Macken, C. A., Hayden, F. G. & Perelson, A. S. Kinetics of influenza A virus infection in humans. J. Virol. 80, 7590–7599 (2006).

38. Pauly, M. D., Procario, M. C. & Lauring, A. S. A novel twelve class fluctuation test reveals higher than expected mutation rates for influenza A viruses. Elife 6, e26437 (2017).

39. Imai, M. et al. Highly pathogenic avian H5N1 influenza A virus replication in ex vivo cultures of bovine mammary gland and teat tissues. Emerg. Microbes Infect. 14, 2450029 (2025).

40. Maines, T. R. et al. Effect of receptor binding domain mutations on receptor binding and transmissibility of avian influenza H5N1 viruses. Virology 413, 139–147 (2011).

41. Herfst, S. et al. Airborne transmission of influenza A/H5N1 virus between ferrets. Science 336, 1534–1541 (2012).

42. Jia, N. et al. Glycomic characterization of respiratory tract tissues of ferrets: implications for its use in influenza virus infection studies. J. Biol. Chem. 289, 28489–28504 (2014).

43. Jayaraman, A. et al. Decoding the distribution of glycan receptors for human-adapted influenza A viruses in ferret respiratory tract. PLoS One 7, e27517 (2012).

44. Fukami, T. Integrating niches, species pools, and priority effects. Annu. Rev. Ecol. Evol. Syst. 46, 1–23 (2015).

45. Stroud, J. T. et al. Priority effects transcend scales and disciplines in biology. Trends Ecol. Evol. 39, 677–688 (2024).

46. Urban, M. C. & De Meester, L. Community monopolization: local adaptation enhances priority effects in an evolving metacommunity. Proc. Biol. Sci. 276, 4129–4138 (2009).

47. Amato, K. A. et al. Influenza A virus undergoes compartmentalized replication in vivo dominated by stochastic bottlenecks. Nat. Commun. 13, 3416 (2022).

48. Castellano, S. et al. IVar, an interpretation-oriented tool to manage the update and revision of variant annotation and classification. Genes (Basel) 12, 384 (2021).

49. Nelson, C. W., Moncla, L. H. & Hughes, A. L. SNPGenie: estimating evolutionary parameters to detect natural selection using pooled next-generation sequencing data. Bioinformatics 31, 3709–3711 (2015).

