## Supplementary Information for "Evolutionary constraints limit additional mammalian adaptation of bovine-derived H5N1 influenza viruses in ferrets"

This file contains:

Supplementary Figs. 1–6

Supplementary Table 1. Sequencing primers

Descriptions of Supplementary Data 1–4

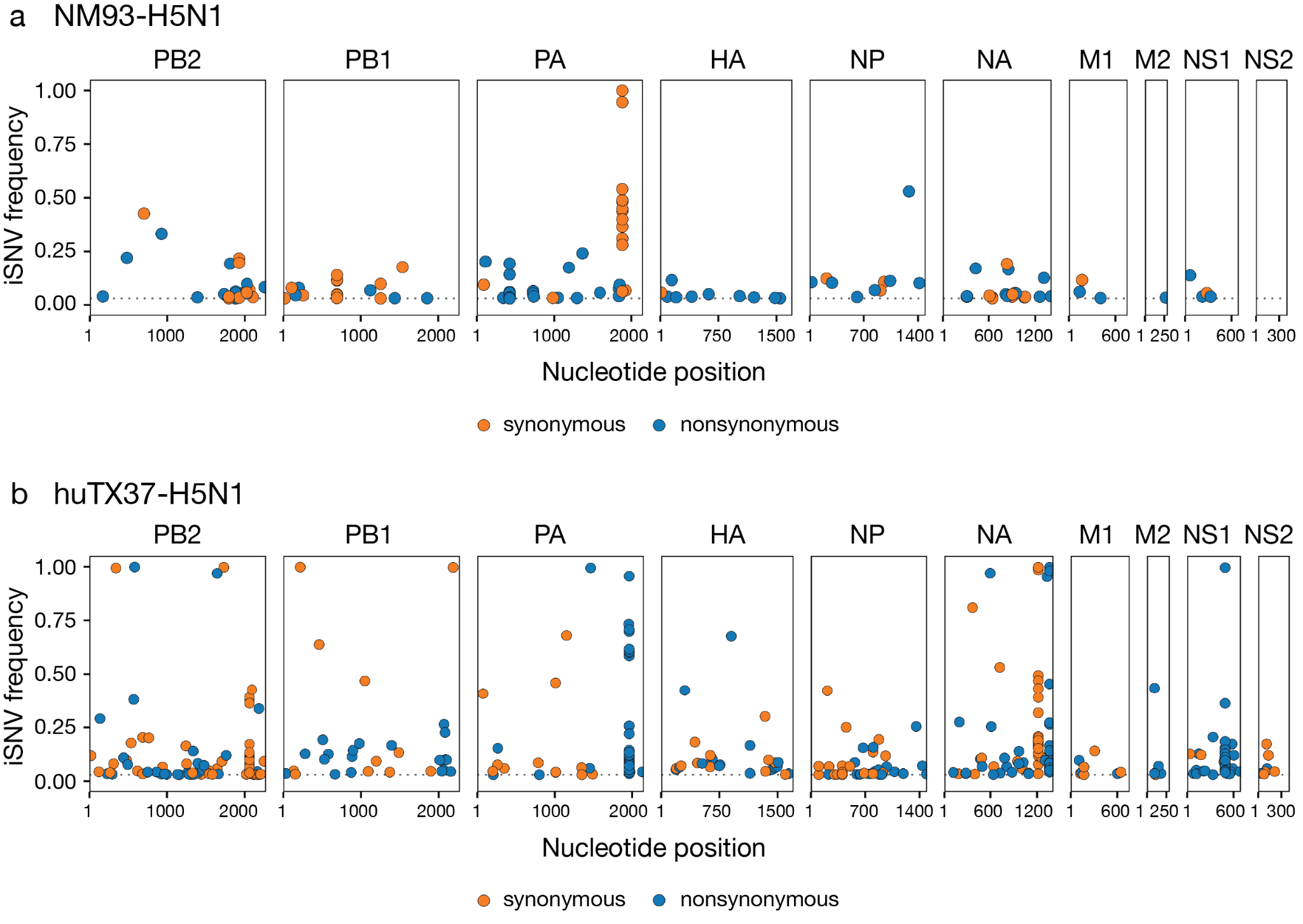

**Supplementary Figure 1.** Distribution of intrahost single-nucleotide variants (iSNVs) across the viral genome by genomic position and allele frequency. (a) NM93-H5N1. (b) huTX37H5N1. Non-synonymous variants are shown in blue, and synonymous variants are shown in orange.

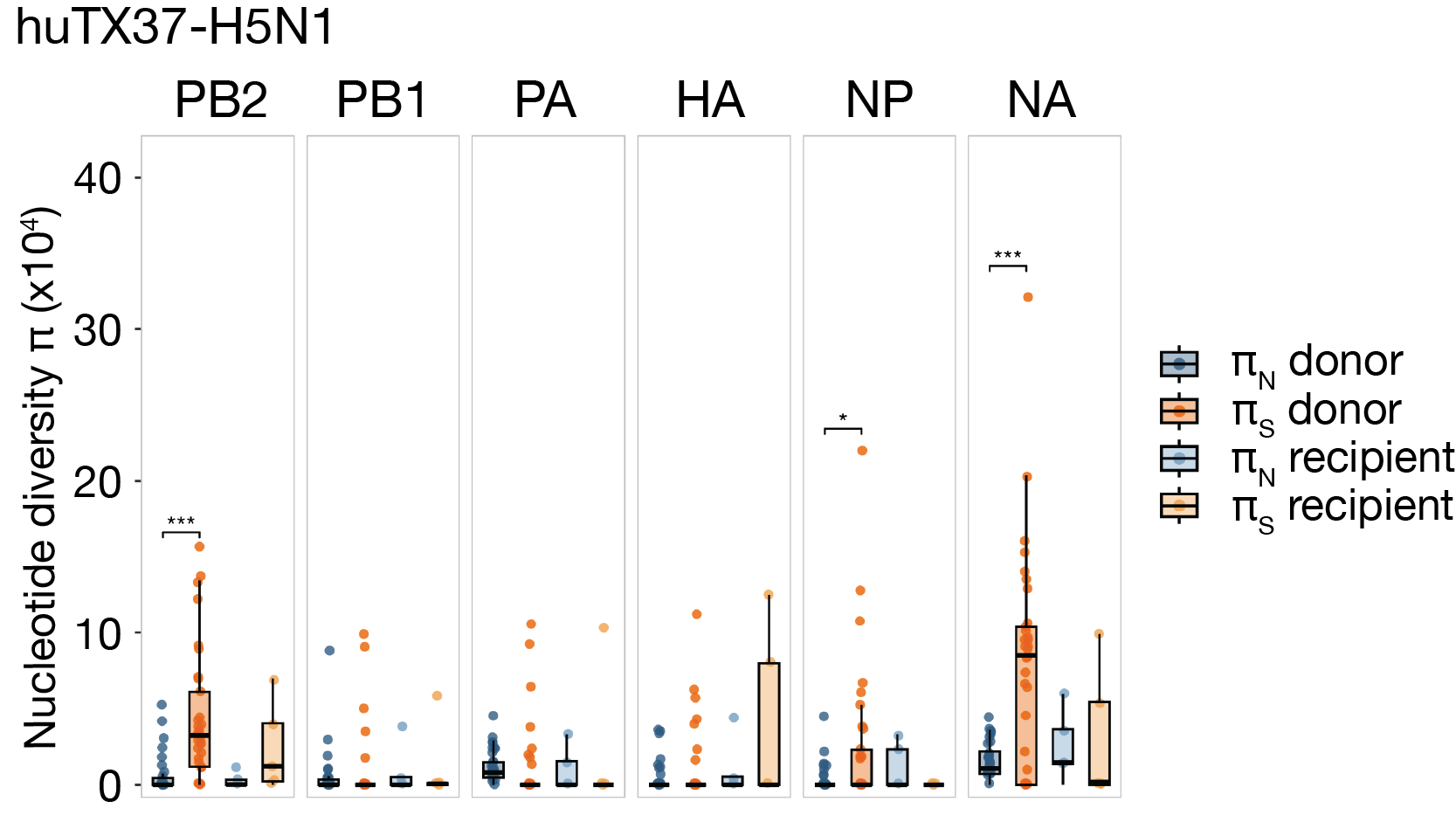

**Supplementary Figure 2.** πN and πS nucleotide diversity is plotted for each gene segment for ferrets infected with huTx37-H5N1 stratified by donor and recipient ferrets for all timepoints. Wilcoxon signed-rank tests were used to test the null hypothesis that the median difference between π values. Bonferroni correction was applied. Asterisks indicate significance: ***** p < 0.05; ****** p < 0.01; ******* p < 0.001.

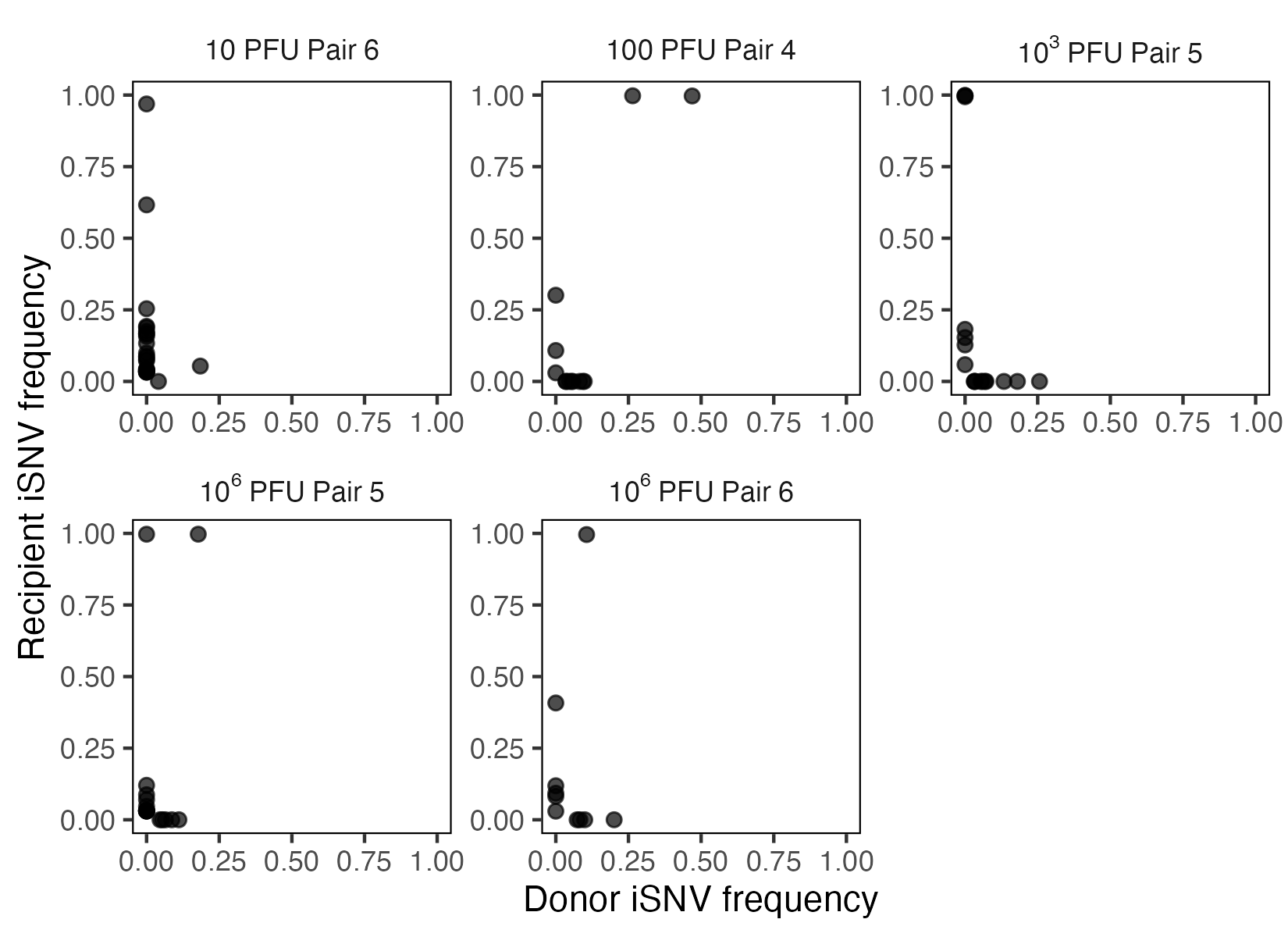

**Supplementary Figure 3.** iSNV frequency in donor versus recipient ferrets in successful transmission pairs for huTX37-H5N1. Latest donor day and earliest recipient time points shown.

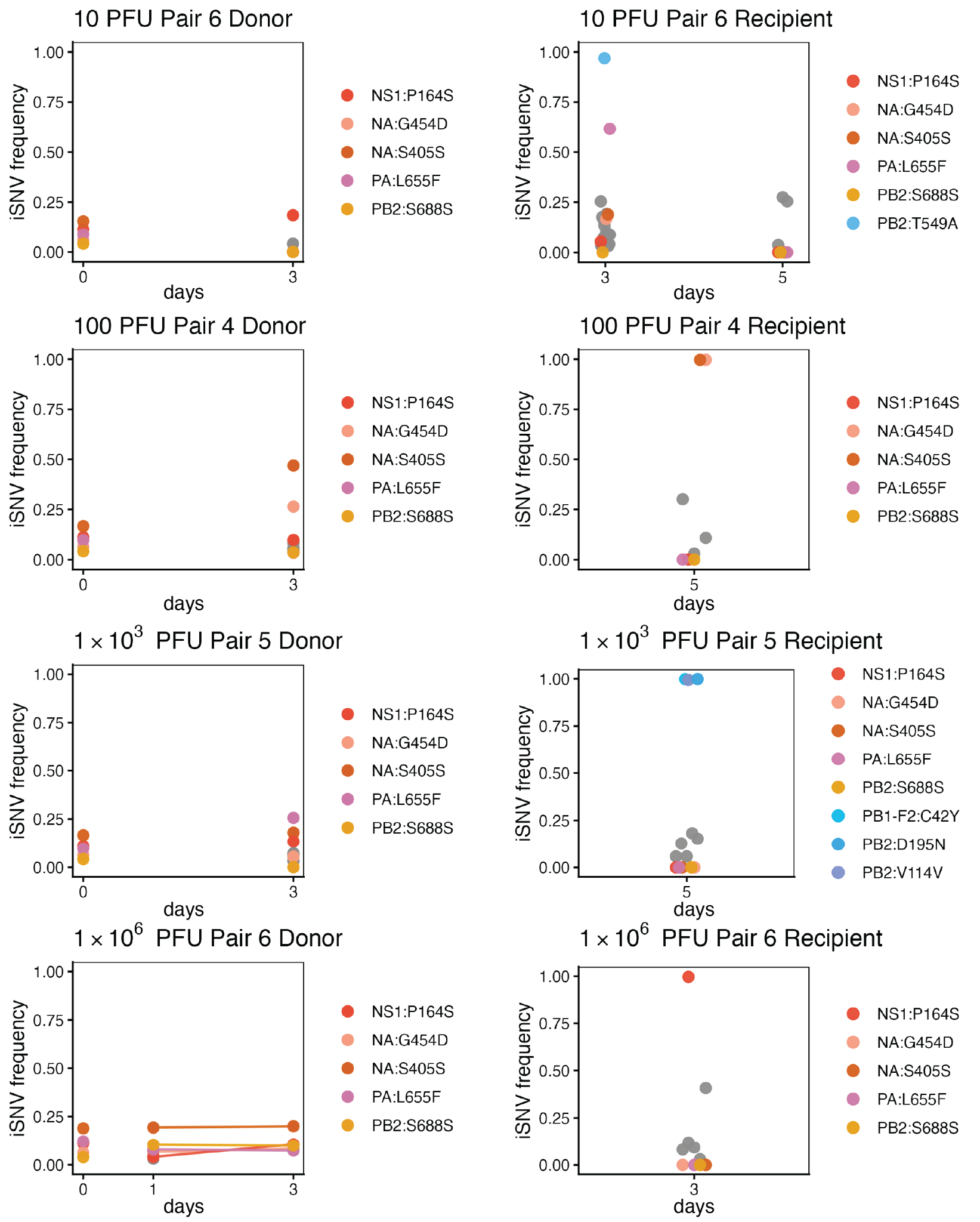

**Supplementary Figure 5.**  iSNV frequency trajectory dynamics across remaining successful transmission pairs for huTX37-H5N1. iSNV trajectories in stock virus (labeled as day 0), donor, and recipient ferret. iSNVs found in stock are colored by red/orange, those that are present in more than one animal or timepoint or present at >50% frequency are in blue/green. Other iSNV present at a single timepoint at <50% frequency are displayed in gray.

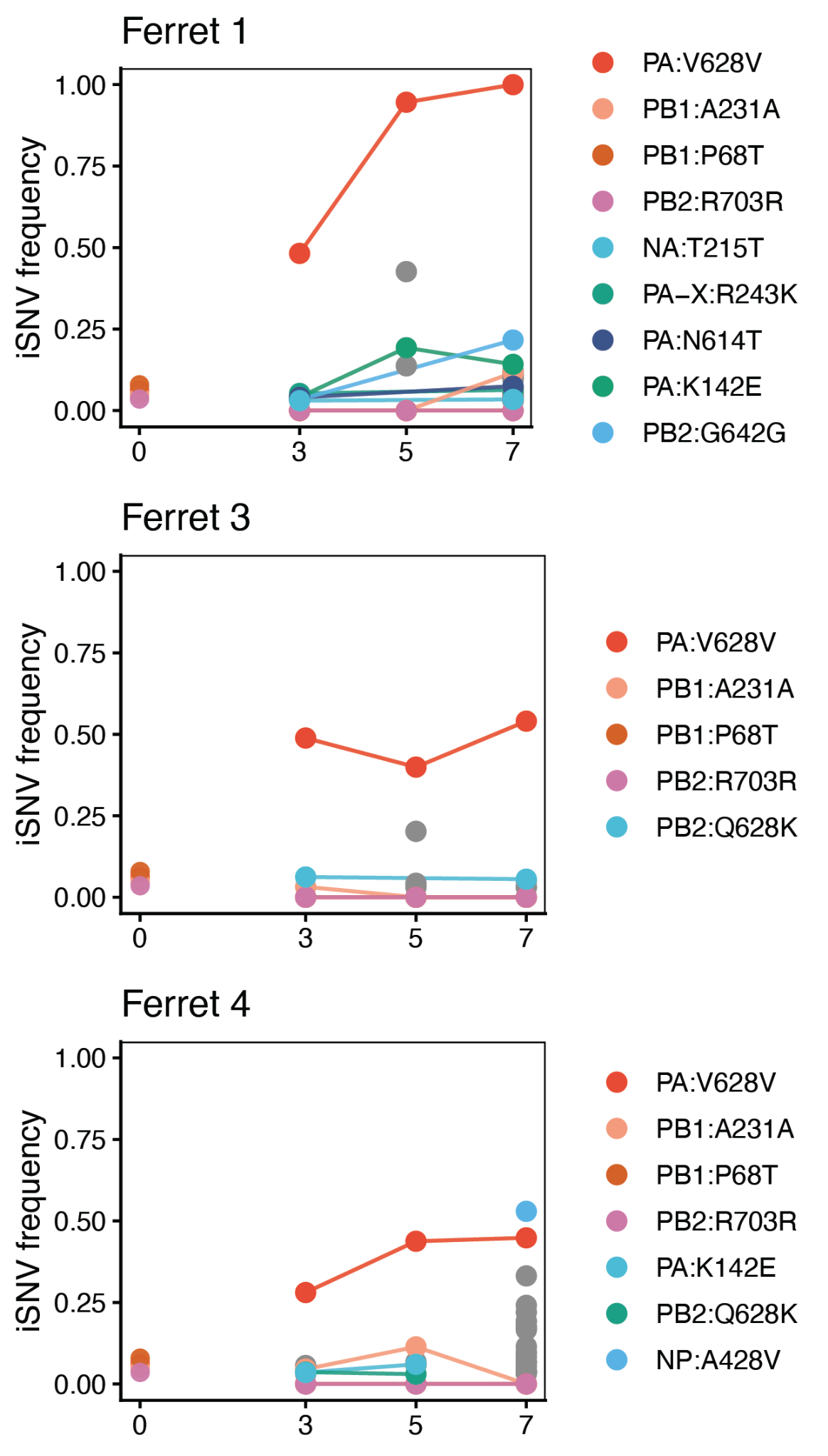

**Supplementary Figure 6.**  iSNV frequency trajectory dynamics across remaining inoculated donors for NM93-H5N1. iSNV trajectories in stock virus (labeled as day 0), and days post infection in the donor ferrets. iSNVs found in stock are colored by red/orange, those that are present in more than one animal or timepoint or present at >50% frequency are in blue/green. Other iSNV present at a single timepoint at <50% frequency are displayed in gray.

**Supplementary Table 1.** Forward and reverse primers used in viral sequencing for each genome segment for NM93-H5N1.

| **Primer** | **Sequence** |
| --- | --- |
| **PB2-F** | **5'-TCGTCGGCAGCGTCAGATGTGTATAAGAGACAGAGCGAAAGCAGGTCAAATATATTCA-3'** |
| **PB2-R** | **5'-GTCTCGTGGGCTCGGAGATGTGTATAAGAGACAGAGTAGAAACAAGGTCGTTTTTAAACAATTC-3'** |
| **PB1-F** | **5'-TCGTCGGCAGCGTCAGATGTGTATAAGAGACAGAGCGAAAGCAGGCAAACCATTT-3'** |
| **PB1-R** | **5'-GTCTCGTGGGCTCGGAGATGTGTATAAGAGACAGAGTAGAAACAAGGCATTTTTTCATGAAGGA-3'** |
| **PA-F** | **5'-TCGTCGGCAGCGTCAGATGTGTATAAGAGACAGAGCAAAAGCAGGTACTGATTCAAAATG-3** |
| **PA-R** | **5'-GTCTCGTGGGCTCGGAGATGTGTATAAGAGACAGAGTAGAAACAAGGTACTTTTTTGGACAGTA -3'** |
| **HA-F** | **5'-TCGTCGGCAGCGTCAGATGTGTATAAGAGACAGAGCAAAAGCAGGGGTTCACT-3'** |
| **HA-R** | **5'-GTCTCGTGGGCTCGGAGATGTGTATAAGAGACAGAGTAGAAACAAGGGTGTTTTTAACTACAAT-3'** |
| **NP-F** | **5'-TCGTCGGCAGCGTCAGATGTGTATAAGAGACAGAGCAAAAGCAGGGTAGATAATCA-3'** |
| **NP-R** | **5'-GTCTCGTGGGCTCGGAGATGTGTATAAGAGACAGAGTAGAAACAAGGGTATTTTTCTTTAATTG-3'** |
| **NA-F** | **5'-TCGTCGGCAGCGTCAGATGTGTATAAGAGACAGAGCAAAAGCAGGAGTTCAAAATGAA-3'** |
| **NA-R** | **5'-GTCTCGTGGGCTCGGAGATGTGTATAAGAGACAGAGTAGAAACAAGGAGTTTTTTGAACAAACT-3'** |
| **M-F** | **5'-TCGTCGGCAGCGTCAGATGTGTATAAGAGACAGAGCAAAAGCAGGTAGATATTGAAAGATGA-3'** |
| **M-R** | **5'-GTCTCGTGGGCTCGGAGATGTGTATAAGAGACAGAGTAGAAACAAGGTAGTTTTTTACTCCAGC-3'** |
| **NS-F** | **5'-TCGTCGGCAGCGTCAGATGTGTATAAGAGACAGAGCAAAAGCAGGGTGACAA-3'** |
| **NS-R** | **5'-GTCTCGTGGGCTCGGAGATGTGTATAAGAGACAGAGTAGAAACAAGGGTGTTTTTTATTATTAA-3'** |

**Supplementary Data 1. NM93-H5N1 iSNV table.** Tabular CSV file containing all iSNVs detected in NM93-H5N1 samples at ≥3% frequency and ≥200× coverage.

**Supplementary Data 2. huTX37-H5N1 iSNV table.** Tabular CSV file containing all iSNVs detected in huTX37-H5N1 samples at ≥3% frequency and ≥200× coverage.

**Supplementary Data 3. Sample identifiers, associated metadata, and SRA accession numbers.**

**Supplementary Data 4. NM93-H5N1 GenBank consensus sequence accessions.** Tabular CSV file containing GenBank accession numbers for NM93-H5N1 consensus sequences.
